# Classification and clustering of RNA crosslink-ligation data reveal complex structures and homodimers

**DOI:** 10.1101/2021.08.01.454689

**Authors:** Minjie Zhang, Irena T. Fischer-Hwang, Kongpan Li, Jianhui Bai, Jian-Fu Chen, Tsachy Weissman, James Y. Zou, Zhipeng Lu

## Abstract

The recent development and application of methods based on the general principle of “crosslinking and proximity ligation” (crosslink-ligation) are revolutionizing RNA structure studies in living cells. However, extracting structure information from such data presents unique challenges. Here we introduce a set of computational tools for the systematic analysis of data from a wide variety of cross-link-ligation methods, specifically focusing on read mapping, alignment classification and clustering. We design a new strategy to map short reads with irregular gaps at high sensitivity and specificity. Analysis of previously published data reveals distinct properties and bias caused by the crosslinking reactions. We perform rigorous and exhaustive classification of alignments and discover 8 types of arrangements that provide distinct information on RNA structures and interactions. To deconvolve the dense and intertwined gapped alignments, we develop a network/graph-based tool CRSSANT (Crosslinked RNA Secondary Structure Analysis using Network Techniques), which enables clustering of gapped alignments and discovery of new alternative and dynamic conformations. We discover that multiple crosslinking and ligation events can occur on the same RNA, generating multi-segment alignments to report complex high level RNA structures and multi-RNA interactions. We find that alignments with overlapped segments are produced from potential homodimers and develop a new method for their de novo identification. Analysis of overlapping alignments revealed potential new homodimers in cellular noncoding RNAs and RNA virus genomes in the Picornaviridae family. Together, this suite of computational tools enables rapid and efficient analysis of RNA structure and interaction data in living cells.

## Introduction

RNA forms complex structures and interactions to execute a wide variety of biological functions. The information-structure duality of RNA underlies its pioneering position in the early evolution of life on earth (Higgs and Lehman 2015). In addition to acting as the messenger between the genetic blueprint and the protein products, structured RNA molecules play extensive roles in scaffolding, regulation, and catalysis in the modern RNA world (Cech and Steitz 2014; Guil and Esteller 2015). Given the importance of this biopolymer, many methods have been developed to determine its structures. Predicting the basepairing of nucleotides, or RNA secondary structure, has long been the goal of algorithms that calculate minimal free energy conformations or exhaustively search for conserved structural motifs in multiple alignments of nucleotide sequences (Gutell 1993; Mathews 2006). Various energy and statistics based computational tools have been developed to predict RNA 3D structures (Das et al. 2010; Weinreb et al. 2016; Miao and Westhof 2017; Sun et al. 2017). On the other hand, classical physical methods, such as X-ray crystallography, nuclear magnetic resonance spectroscopy and cryo-EM have made significant progress in recent years towards solving more complex 3D structures of RNAs and their complexes (Batey et al. 1999; Bai et al. 2015).

In the last few decades, a host of chemical methods were invented to probe the flexibility and accessibility of individual nucleotides, which are indicative of their structural context (Weeks 2010; Lu and Chang 2016; Velema and Kool 2020). These methods typically yield indirect 1-dimension information that assist secondary and tertiary structure prediction. More recently, several crosslinking-based methods, including CLASH, hiCLIP, PARIS, LIGR-seq, SPLASH, fRIP and COMRADES, have been advanced to provide direct physical evidence for spatial proximity among RNA fragments (Kudla et al. 2011; Helwak et al. 2013; Sugimoto et al. 2015; Aw et al. 2016; Hendrickson et al. 2016; Lu et al. 2016; Nguyen et al. 2016; Sharma et al. 2016; Lu et al. 2018; Ziv et al. 2018; Lu et al. 2020; Zhang et al. 2021). These methods employ a variety of crosslinkers, such as psoralens that only react with staggered uridines and cytidines in opposing strands, UV that reacts with both proteins and RNAs in direct contacts, and formaldehyde that crosslinks all types of primary amine-containing molecules that are close to each other (Lu and Chang 2018). After crosslinking, and purification/enrichment, covalently attached RNA fragments are ligated and sequenced in high throughput, yielding hybrid reads, where each segment comes from a distinct region in an RNA, or from entirely different RNA molecules. In the simplest form, the crosslink-ligation experiments reveal RNA hetero-duplexes on a transcriptome wide scale (**Fig. 1A**). In reality, hetero duplexes with two arms are not the only form of structures in RNA structures and interactions, other types of complex arrangements are also common and critical for the formation of high-level structures.

**Figure 1.**
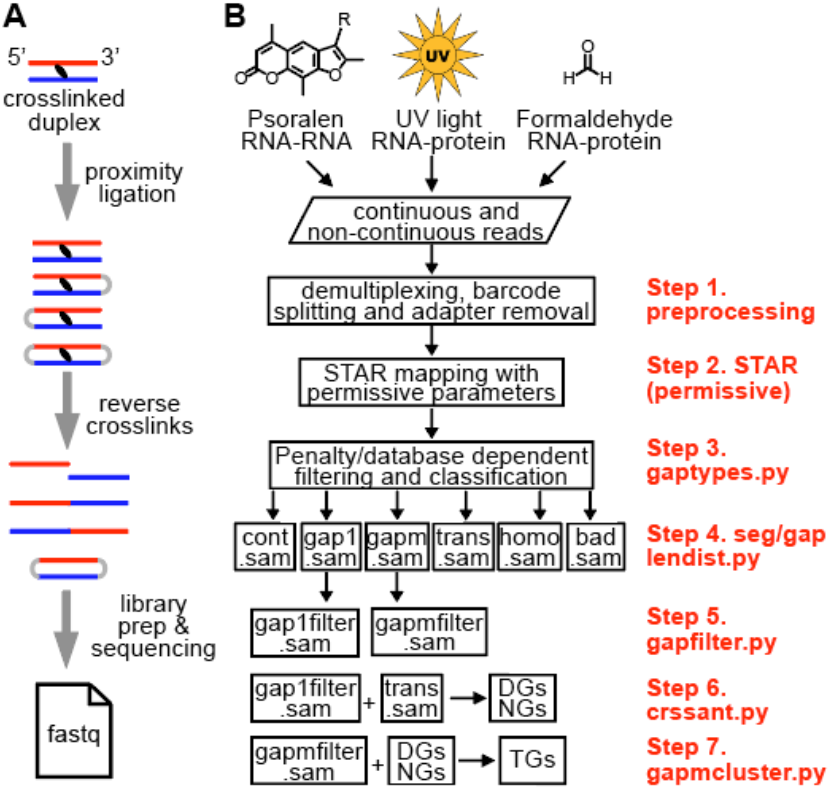
Overview of RNA crosslink-ligation experiments and analysis pipeline. (A) Outline of a typical crosslink-ligation experiment leading to fastq output files. The proximity ligation of crosslinked duplexes can produce both forward and backward arrangements. Circularized RNAs are rare and lost during library preparation, because they cannot be ligated to adaptors. Similarly, concurrent crosslinking at multiple locations and subsequent ligation of them produce multi-gapped reads (gapm in panel B). (B) Several different types of crosslinking methods, such as psoralen, UV and formaldehyde, together with proximity ligation produces non-continuous reads that can be used to determine RNA structures. Newly developed computational tools and optimized parameters are listed on the right in 7 steps. Reads are preprocessed (step 1) and mapped to genome references using optimized STAR parameters (step 2), filtered and classified to 6 alignment types (step 3), including continuous (cont.sam in SAM format), one-gap (gap1), multi-gap (gapm), trans interactions (trans), homotypic interactions (homodimers, or homo), and miscellaneous bad alignments (bad). After quality control and removal of splicing events and artifacts (steps 4-5), each of these alignment types are further processed to extract information for duplexes (step 6), high-level structures (step 7), and RNA homodimers (homo.sam).

First, within the same molecule, high level structures include extended helices with various internal loops, multi-helix junctions, pseudoknots, and even triple helices. In the past 50 years, in vitro studies have shed light on the exquisite folding a number of RNAs and their complexes, such as the ribosome, RNase P, RMRP, telomerase, mascRNA, viral IRES elements, each employing unique combinations of the aforementioned high-level structures (Wilusz et al. 2012; Anger et al. 2013; Quade et al. 2015; Zhang et al. 2017; Wu et al. 2018; Kastner et al. 2019; Yan et al. 2019). However, direct in vivo observation of these complex structures and interactions has been more difficult, despite their demonstrated functional significance in well studied examples.

Second, between different RNA molecules, homodimers are also possible besides heterodimers, yet very few RNA homodimers have been studied, despite the high stability of the base pairing interactions (Bou-Nader and Zhang 2020). Examples have been reported in a variety of contexts, including viral RNA genomes, such as HIV, HCV, coronaviruses and bacteriophages (Clever et al. 2002; Shetty et al. 2010; Ishimaru et al. 2013; Dubois et al. 2018), ribozymes and riboswitches (Bou-Nader and Zhang 2020), mRNAs (Wagner et al. 2004; Jambor et al. 2011; Little et al. 2015; Trcek et al. 2015), trinucleotide/ hexanucleotide repeats (Ciesiolka et al. 2017; Jain and Vale 2017), tRNA mutant and fragment dimers/tetramers (Wittenhagen and Kelley 2002; Roy et al. 2005; Lyons et al. 2017; Tosar et al. 2018). In vitro, synthetic RNAs have also been made to dimerize or multimerize to prepare nanomachines (Severcan et al. 2009; Geary et al. 2011). These homodimers play important roles in virus genome packaging, stress response, translational regulation, liquid-liquid phase separation and human genetic diseases. Again, de novo identification of homodimers remains challenging.

Despite the rapid progress in crosslink-ligation experimental techniques, there are three major challenges in the data analysis. First, the random RNA fragmentation by RNases or divalent cations and subsequent proximity ligation generates short reads with irregular gaps. Longer reads can be mapped to references with higher accuracy, but the resolution of secondary structure models is lower. Shorter reads increase the model resolution, but mapping accuracy is lower. A number of short read mappers have been developed with the ability to handle gaps. For example, Bowtie2 applies affine penalty to gaps, which discourages gap opening and extension (Langmead and Salzberg 2012). Multi-step mapping protocols based on Bowtie2 reduces sensitivity for shorter segments that cannot be mapped uniquely to the genomes (Travis et al. 2014; Sharma et al. 2016). STAR can inherently map non-continuous reads, but the parameters were optimized for the identification of splicing and gene fusion events (Dobin et al. 2013; Haas et al. 2017), and performances were suboptimal on crosslink-ligation data (Aw et al. 2016; Lu et al. 2016; Ziv et al. 2018). For example, splice junctions have unique sequence consensus to facilitate opening of gaps and assignment of extension penalty; in addition, splice junction databases can be used to help mapping, reducing the unnecessary penalty and increasing the efficiency. Non-continuous reads from crosslink-ligation experiments, however, are far more random in gap sequence and length, making it difficult to determine the appropriate penalty. To solve this problem, we systematically optimized the STAR parameters in this study, and designed a set of filtering criteria that significantly improved the sensitivity and specificity of mapping short reads with irregular gaps.

Second, in addition to simple duplexes, the complex crosslinking and proximity ligation reactions produce many different types of reads/alignments that remain poorly characterized. Our exhaustive classification uncovered 8 categories of alignments, which we rearrange and combine to 5 distinct types, including continuous (cont for short), two-segment (1 gap, or gap1), multi-segment (>1 gaps or >2 segments, gapm), homodimers (overlapped segments, homo), and trans interactions (two segments on different strands or chromosomes). Each type of non-continuous alignments reveals distinct new structures and interactions, especially composite structures and homodimers. The rearrangements also enable the visualization and of complex alignments in genome browsers and facilitate intuitive understanding of their corresponding structures.

Third, densely packed non-continuous alignments are difficult to deconvolve into distinct groups that support individual RNA duplexes, because most RNA duplexes are very short and close to each other. This is further complicated by the multitude of alternative/dynamic conformations, where one RNA region can base pair with multiple rother egions. To resolve the complex structure conformations encoded in non-continuous alignments, we developed a method to cluster alignments based on a network representation, termed CRSSANT. Alignments are assigned to duplex groups (DGs) based on segment overlap ratios, and then DGs can be used to constrain secondary structure modeling. This new method is automatic and separates alternative conformations from each other. Using DGs as the foundation, we further developed a method to build tri-segment groups (TGs) that reveal high-level structures and interactions among RNAs.

Using these newly developed tools, we systematically characterized published crosslink-ligation methods, revealing their basic properties and bias. For example, we noticed that psoralen monoadducts lead reverse transcription errors and over-representation of uridine deletions in some of these crosslink-ligation methods. Our classification and clustering of various types of alignments led to the discovery of high-level structures and interactions and RNA homodimers in various cellular and viral RNAs. Together this suite of tools greatly expanded the capabilities of crosslink-ligation experimental methods.

## Results

### Overview of the computational pipeline

In general, crosslink-ligation experiments produce several types of reads, including continuous and non-continuous, where the continuous reads could be due to failed crosslinking or ligation, while non-continuous ones may contain 2 or more segments (**Fig. 1A**, showing 2-segment reads as examples). To extract all possible types of structures from crosslink-ligation data, we established a general strategy that is applicable to different types of experimental strategies, including, but not limited to, psoralen, UV or formaldehyde crosslinking (Kudla et al. 2011; Sugimoto et al. 2015; Aw et al. 2016; Hendrickson et al. 2016; Lu et al. 2016; Sharma et al. 2016; Van Nostrand et al. 2016; Ziv et al. 2018) (**Fig. 1B**). The sequenced reads are first processed to remove adapters, barcodes, and demultiplexed using common tools (step 1, e.g. FASTX and Trimmomatic) (Bolger et al. 2014). Processed reads are mapped to genome references using STAR (Dobin et al. 2013) and a set of parameters that we specifically optimized for non-continuous reads (step 2). The optimized STAR method and subsequent filtering maximize the sensitivity and specificity in the analysis of short segments with irregular gaps. Alignments are filtered to remove low-confidence segments, rearranged, and classified into 6 categories (step 3). In addition to simple RNA duplexes, these different types of alignments provide new information such as high-level structures (multi-gap alignments, or gapm), and RNA homodimers (homotypic interactions, or homo). Segment and gap length distribution and gap nucleotide properties are summarized to serve as quality controls (step 4). Gapped alignments (with 1 or more gaps) are filtered to remove splicing junctions and short gaps that are likely artifacts (step 5). The filtered non-continuous alignments are clustered into duplex groups (DG) and non-overlapping groups (NG, for visualization in genome browsers) (step 6). The alternative conformations (conflicting DGs) suggest the existence of dynamic RNA structures and functions. Multi-gap alignments together with DGs are further clustered into TGs that support more complex structures and interactions (step 7).

### Optimized short read mapping and filtering of crosslink-ligation sequencing data

The first critical step in analyzing crosslink-ligation data is mapping short reads with high sensitivity and specificity. To demonstrate the relevance of read length in structure modeling, we examined RNA duplexes in well-studied structures, the human ribosome and spliceosome (Petrov et al. 2014; Yan et al. 2019) (**Fig. 2A-B, Supplemental Fig. 1A**). We found that ~91% of arms are <= 20nt, and more than 50% of them are <=10nt. Bowtie2 can map parts of reads and the separately mapped segments can be chained to identify the gaps. The multi-step mapping strategy results in low sensitivity since both segments need to be long enough (e.g. >=20nt) for unique mapping. The gap penalty is linear to gap size, making it difficult to accommodate long gaps. STAR considers the multiple segments together when calculating alignment scores. In addition, gap penalty calculation is more flexible, making it possible to retain short segments. Several previous studies used minimally modified STAR parameters (Ramani et al. 2015; Aw et al. 2016), while others used Bowtie2 and additional post-processing (Sugimoto et al. 2015; Nguyen et al. 2016; Sharma et al. 2016; Yu et al. 2016) (**Supplemental Table 1**).

**Figure 2.**
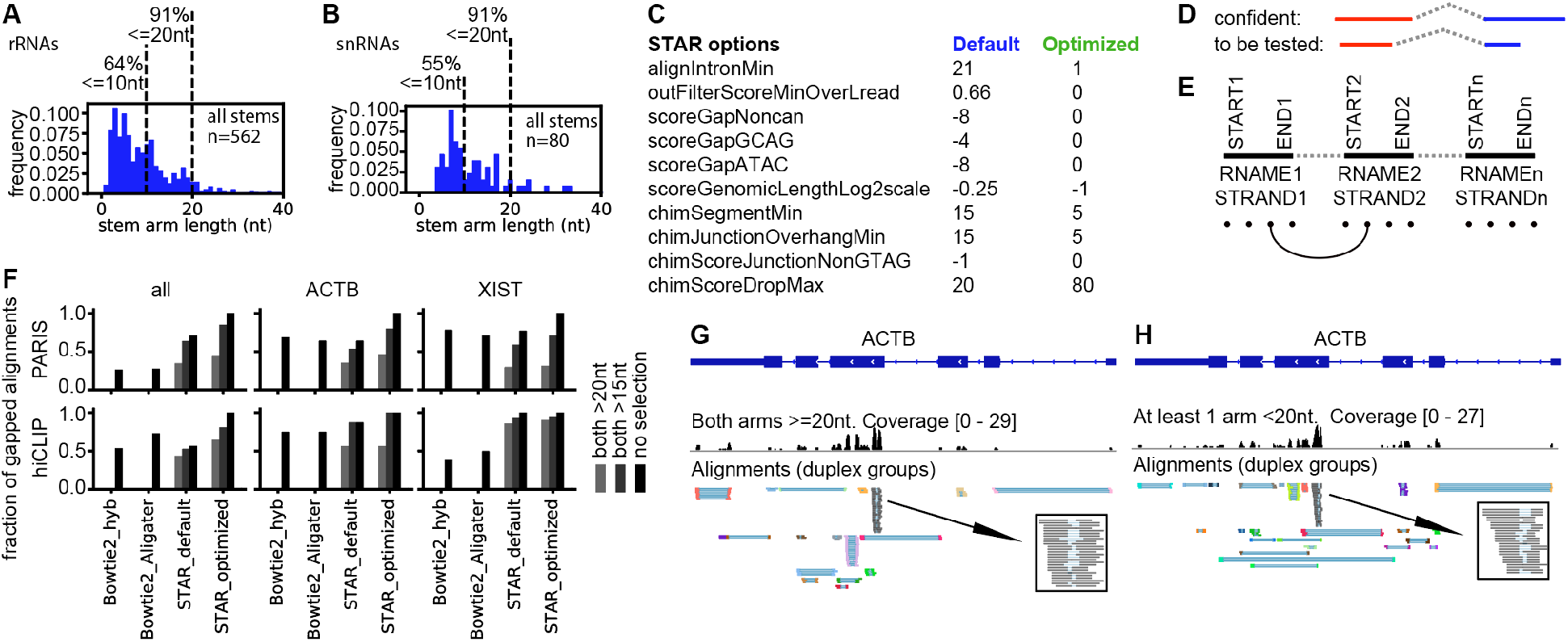
Optim ization of short read mapping from crosslink-ligation experiments. (A-B) RNA stems were extracted from the human cytoplasmic and mitochondrial ribosome and spliceosome crystal or cryo-EM structures. The following RNAs are included: 12S, 16S, 5S, 5.8S 18S, 28S, U1, U2, U4, U6, U5, U11, U12, U4atac and U6atac. (C) List of critical STAR parameters that are optimized to map non-contlnuous reads. The default value for chlmSegmentMin Is unset, whereas setting this value to any positive integer trlg-gers chimeric alignments. The recommended value of 15 is used here as the “default” for comparison. (D-F) Strategies for filtering alignments after STAR mapping. (D) Confident alignments: all segments or arms are uniquely mapped to the genome. Alignments with shorter segments that cannot be mapped uniquely are to be tested against confident ones. (E) Filtering method for the less confident alignments: all arms of the confident alignments are built into a database of connections between segments, in 5 nucleotide intervals (dots shown at the bottom). The connection database consists of reference name (RNAME), strand (STRAND) and coordinates between start and end (START, END). Then the less confident alignments are tested against this database. (F) Gap1 (one gap, i.e. two segments) alignments in PARIS and hiCLIP data were recovered by various mapping methods and segment-length selections. Fractions for the highest-performing method (STAR_optimized) are set to 1. For STAR analysis, sequencing reads were mapped to the genome (hg38 primary); then alignments were filtered and classified to 6 categories using gaptypes.py. The gap1 alignments were filtered to remove short gaps and splicing alignments (gapfilter.py). Primary alignments were extracted from all alignments and used for analysis. For Bowtie2 mapping, previously reported parameters (hyb and Aligater) were used. Unique alignments with deletions (D in SAM CIGAR string) were extracted and alignments were converted to join the multiple segments (bowtie2chlm.py). Then the alignments were classified using gaptypes.py. The gap1 alignments were filtered to remove short gaps and splicing alignments (gapfilter.py). The selection of alignments with both arms > 15nt or 20nt mimics the mapping and chaining strategy in previous studies that employ Bowtie2 (hyb and Aligater). (G-H) Alignments in the ACTB mRNA from PARIS data in HEK cells were separated to ones where both arms (or segments) are at least 20nt (G), or at least one arm is shorter than 20nt (H). The inset boxes show DGs that support the same duplex regardless of segment length.

Here we used STAR to develop a new strategy to identify gapped reads with high sensitivity and specificity. In principle, STAR searches for maximal mappable prefixes (MMP) sequentially from fragments of the sequencing read, starting from the first base (Dobin et al. 2013). Here junctions are detected naturally during the iterative search process and all types of junctions or gaps are included. After all MMPs are detected, they are clustered, stitched and scored in the second step. All seeds that are within the user defined genomic windows are stitched (default: winBin = 2A16, and window = 9*winBin = 589824). The principle for our new strategy is that we allow mapping of short fragments by (1) removing penalty for gap opening (scoreGap* parameters in STAR), (2) reducing penalty for gap extension (scoreGenomicLengthLog2scale in STAR), and (3) allowing chimeric alignments with short fragments (**Figure 2C, Supplemental Table 2**. Methods and Supplementary Material). This combination of new parameters effectively treats all gaps like splicing junctions. After mapping, the alignments are filtered based on (1) segment length, and (2) overlap of less-confident shorter segments with confident longer ones (**Fig. 2D-E**). If shorter segments are close to long segments, they are very likely to be unique and bona fide, even though their presence in the entire genome is not unique. If shorter segments overlap longer ones, they are also considered confident. For example, an alignment with CIGAR string 20M30N10M (20nt match, 30nt gap and 10nt match) is likely to be real, because the 10M segment is very close to the 20M segment. The permissive STAR parameters and the filtering enable recovery of shorter fragments.

To systematically evaluate published crosslink-ligation methods and the new mapping strategy, we processed data using uniform procedures (**Supplemental Fig. 1B**). After mapping using the optimized STAR parameters, we classified and rearranged alignments into 6 categories (see details later) and removed spliced alignments. The mapped segments and gaps follow a wide range of distributions (**Supplemental Fig. 2C-E**). PARIS data have a median segment size of 23nt, followed by hiCLIP at 31nt. For PARIS, ~95% of the segments are shorter than 40nt, and a significant portion of them, ~13.8%, at or below 15nt. Other psoralen crosslinking data have median segment sizes above 43nt, about twice the size of PARIS data. Surprisingly, we found that 1-2nt gaps are present in a significant portion of all non-continuous alignments (**Supplemental Fig. 2D-E**). In one SPLASH dataset, 1-2nt gaps are present in 72% alignments, whereas only 9-18% gaps in PARIS are 1-2nt. To determine whether the short gaps are artifacts, we analyzed nucleotide frequencies in the gaps. For data with more 1-2nt gaps, uridine is significantly over-represented (**Supplemental Fig. 2F-G**). SPLASH and COMRADES have significantly higher bias than PARIS and LIGR. SPLASH and COMRADES used biotinylated psoralens for enrichment, where monoadducts at uridines are the dominant products, rather than the crosslinks. In LIGR and PARIS, enrichment crosslinked fragments was achieved using RNase R and 2D gels, respectively, where the monoadduct are much lower. We speculated that such 1-2nt gaps are due to reverse transcription errors on psoralen-uridine monoadducts, therefore we removed them before further analysis. The gap and segment length and composition analysis provided valuable information about the library quality and should server as important guides for future applications and optimizations. Together, PARIS and hiCLIP consistently outperformed other crosslinking methods in the gap and segment length distributions.

Given that PARIS and hiCLIP produced the shortest segments, we used these two to benchmark the optimized STAR mapping and filtering procedure. Specifically, we compared published Bowtie2, default STAR (default in STAR except the activation of the chimeric alignments) and optimized settings (**Fig. 2F**). The optimized mapping and filtering improved recovery of all alignments dramatically over other methods on both PARIS and hiCLIP (black bars, **Fig. 2F**). Among the mapped alignments, roughly 50% of them have both arms > 20nt, which is the commonly used cutoff in multi-step mapping procedures in other studies. From the most stringent condition (default with both arms >20nt), to the most sensitive condition (optimized with no size selection), the mapped non-continuous alignments increased 2.87-fold. To make sure that the differences in mappability is not due to artifacts, we examined the alignments mapped to two RNAs, ACTB and XIST. The results are consistent with global comparison, despite differences in sequence composition and presence of complex repeats in XIST (Lu et al. 2016). As an example, we separated the gapped alignments on the ACTB mRNA into two groups, where both arms are >=20nt (**Fig. 2G**) or at least one arm is <20nt (**Fig. 2H**). Alignments in both length ranges are clustered into duplex groups (DGs) and compared side by side. We found that the DGs are similar between the different size ranges (see the inset boxes). In fact, the shortest segments in the alignments mapped to ACTB mRNA are only 10nt, yet they are still mapped with high confidence. In summary, we showed that different crosslink-ligation protocols produce non-continuous alignments with drastic differences in segment and gap properties. The optimized parameters and postprocessing for STAR mapping significantly improved the recovery of short segments that are most valuable for building high resolution structure models.

### Rearrangement and classification of alignments

The complex reactions of crosslink-ligation produce complex arrangements in each read. Through exhaustive classification, we divided alignments into 8 types (**Fig. 3A**). Non-gapped alignments from non-crosslinked fragments or failed ligations are type 1. Local collinear gapped alignments, within the predefined window (as defined in STAR), are type 2. Alignment segments that are too distant (beyond STAR genomic window), even though collinear, are considered as one type of chimeras (type 3). Types 2 and 3 are artificially separated because STAR treats local and distal segments in different ways. Chimeric alignments also include ones with reversed orders (type 4), two arms overlapped (type 5), located on different strands (type 6) or chromosomes (type 7). Multi-segment alignments are also possible, arising from multiple proximity ligations or a combination of splicing and proximity ligation (type 8). In theory, the CIGAR string in the SAM format can only accommodate collinear arrangements with positive gaps (gap length >0) i.e. types 1-4, but not overlaps (gap length <0) and non-collinear ones. In STAR, types 4-7 and some of 8 are all considered as chimeric and therefore represented by two or more records each.

**Figure 3.**
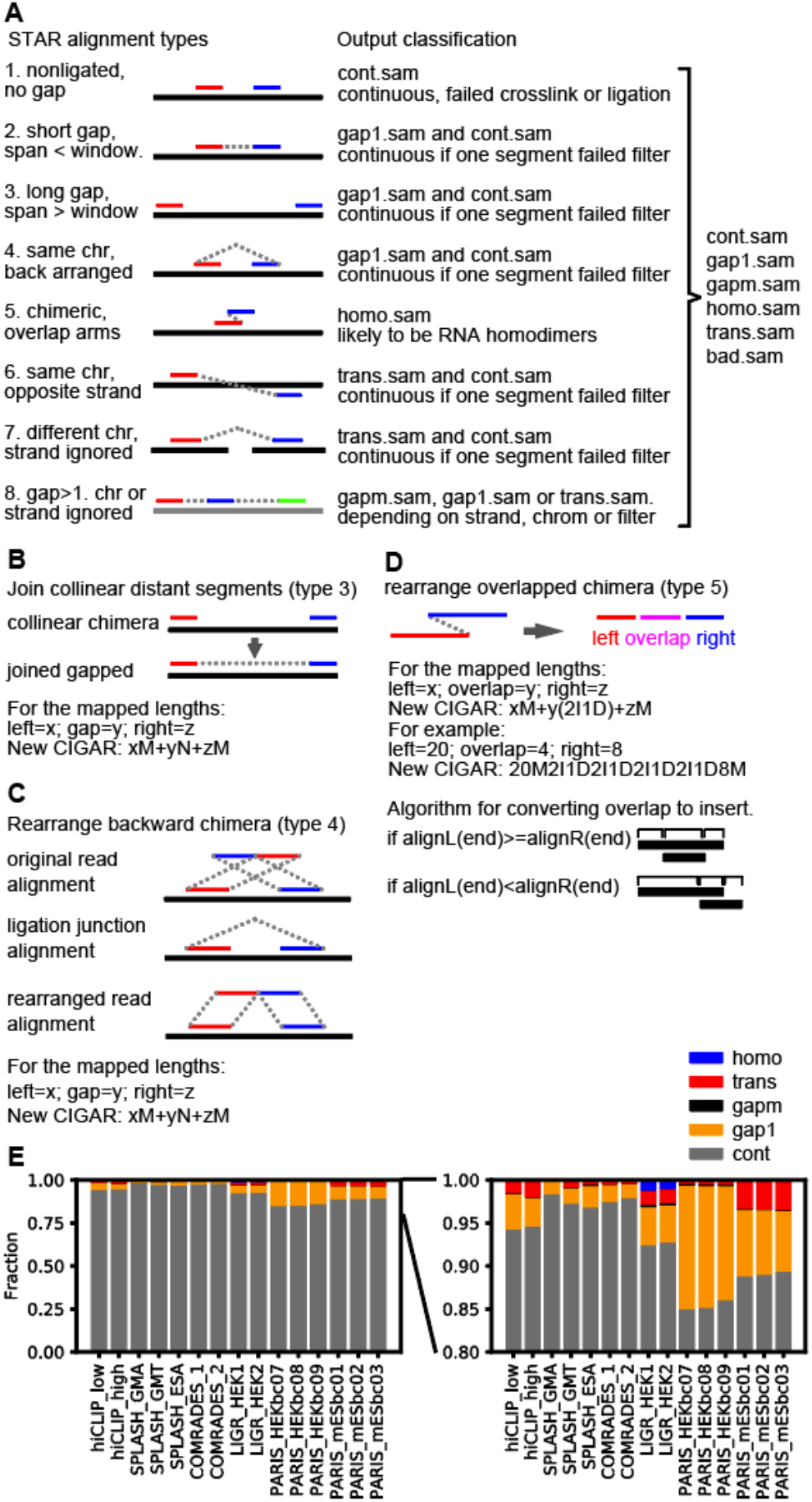
Classification and processing o f alignments from crosslink-li gation experiments. (A) Types of alignments and classification after processing. This diagram presents a unified model for data from all types of crosslink-ligation experiments, and the terms are defined as follows. A read is one piece of sequence from the sequencing machine, and it may have one or multiple alignments to the reference. Segment or arm: part of an alignment with no N’ in the CIGAR substring. Continuous alignments: type 1, with only 1 segment or arm. Gapped: forward arrangement, with 1 or more gaps, including gapl and gapm (types 2 and some of type 8). Chimeric: non-contin uous alignments similar to the definition from the STAR method, including types 3-7 and some of type 8. Non-continuous: including both gapped and chimeric alignments. Homotypic: chimeric alignments where the arms overlap, suggesting RNA homodimers. Trans: segments mapped to different chromosomes or strands (types 6-7 and some of type 8) (B) Diagram for joining collinear distant segments into gapped alignments. The two segments are connected so that the two arms are represented by one record in SAM format. (C) Diagram for rearranging backward chimeric alignments to normal gapped alignments. The 5’ and 3’ arms are switched so that the two segments can be represented by one record in SAM format (D) Diagram for rearranging overlapped chimera The two arms are converted to 3 segments: left overhang, overlap, and right overhang. The new alignment can be reprerented by one record in SAM format (E) Classification of alignments from previously published crosslink-ligation experiments, in which the low abundance categories are magnified on the right.

Even though this exhaustive classification is based on the output from STAR, they are generally applicable to alignments from other types of short read mappers, with minor differences (e.g. local vs. distal gapped in types 2 and 3), and therefore should facilitate more sophisticated studies of RNA structures and interactions. The complex arrangements of the alignments make them difficult to analyze and visualize. Therefore, we developed tools to filter, rearrange and reclassify the 8 types of alignments into 5 types (excluding bad ones), each providing a distinct type of information for inferring RNA structures and interactions (**Fig. 3A**, the right-side classification output, and **3B-E**, see flowchart in **Supplemental Fig. 2** and details in Methods and Supplemental Material). Distant collinear chimeras (type 3) are converted to normal chimeras (gap1, type 2) by joining the two segments (**Fig. 3B**). Backward chimeras (type 4) are converted to normal chimeras (gap1, type 2) by switching the two segments (**Fig. 3C**). Overlapped chimeras (type 5) are converted to homotypic chimeras (homo) by redefining the overlapped part as a combination of insertions and deletions (**Fig. 3D**). After conversion, these types can be processed and visualized as normal gapped alignments. Trans alignments and some of the multi-gap alignments that map to different chromosomes or strands are processed separately (Methods). We applied the alignment, filtering, and rearrangement methods to published datasets (**Fig. 3E**). Each experiment produced variable amounts of alignments in the 5 types (except the bad.sam which are very rare). Even though most homo (overlapping arms) and gam (multi-segment) alignments represent a small percentage of total number of alignments, they are significant given since they reveal important new structures and interactions, and only a few reads/alignments are sufficient to call a specific RNA duplex (see details below).

### Network-based duplex group assembly of single-gapped alignments

Among the 5 rearranged alignment types, gap1, gapm, trans and homo support distinct RNA structures and interactions. To assemble alignments into groups that support individual structures, we developed a method CRSSANT to cluster single gap alignments, including gap1 and trans, to duplex groups (DGs). CRSSANT leverages network analysis techniques -- also frequently referred to as “graph” techniques -- to automate analysis of sequencing reads produced by crosslink-ligation methods. To determine the relationship among alignments, we defined the overlap ratios between any pair of alignments on both arms, o_i_(r_i_,r_2_)/s_i_(r_i_,r_2_) and o_r_(r_i_,r_2_)/s_r_(r_i_,r_2_) (**Fig. 4A**). Then the gap1/trans alignments were converted to a network based on their overlap ratios, and the network is clustered using two alternative approaches, cliques-finding and spectral (**Fig. 4B, Supplemental Fig. 3**, Methods and Supplemental Materials). The clustered subgraphs correspond to individual DGs, each containing highly similar alignments. The clustering produces two types of output, tagged SAM alignments and summary of DG information, which can be used for subsequent visualization and secondary structure prediction.

**Figure 4.**
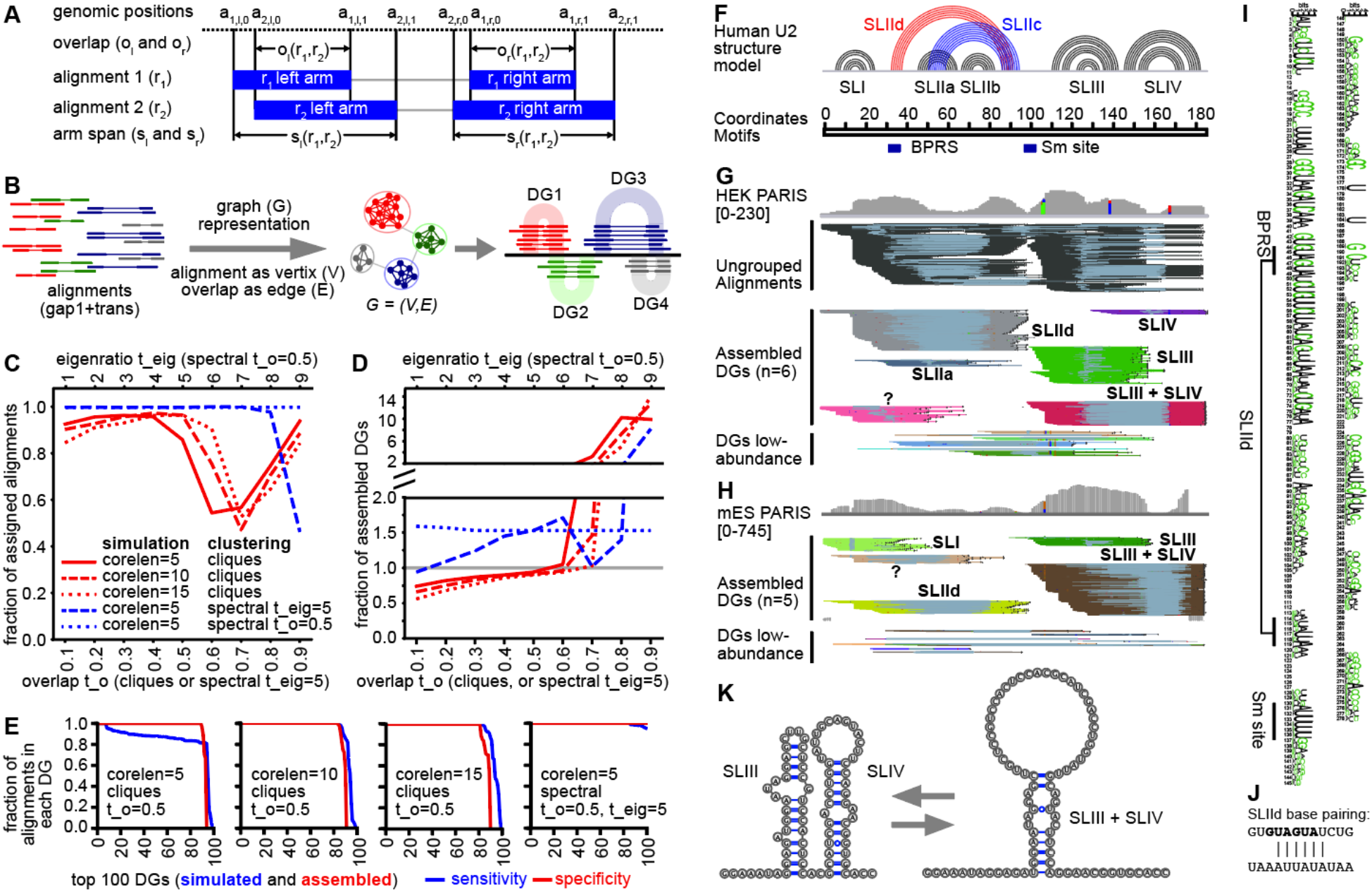
Network/graph-based method for automatic assembly of duplex groups underlying RNA structures and interactions (CRSSANT). (A) Overlap and span calculation for a pair of alignments. Two alignments r_1_ and r_2_ each comprising a left and right arm (solid blue bars), share left and right overlaps o_l_, o_r_, respectively and left and right spans s_l_, s_r_, respectively. The arm start and stop positions of read/allgnment I are represented by the 4-tuple (a_i,l,0_, a_i,l,1_, a_i,r,0_, a_i,r,1_). The two arms can be on the same chromsome and strand (gapl.sam), or different ones (trans.sam). (B) Diagram for network/graph based clustering. All alignments with a single gap (gap1 and trans) are represented as a graph where each alignment Is a vertlx and the relative overlap ratio between the arms Is the edge. Highly connected vertices cluster together forming sub-graphs, corresponding to individual DGs. (C-D) Benchmarking CRSSANT clustering on 100 simulated DGs. All alignments map to chr1:1-1000, and consists of cores 5, 10 or 15nts (corelen=5, 10 or 15), and random extensions on each side between 5 and 15nts. Gaps between the two cores are at least 50nts and at most the length of the chr1:1-1000. Each DG contains between 10 and 100 alignments. The alignments were clustered using cliques or spectral algorithms. For cliques, overlap threshold t_o was varied between 0.1 and 0.9. For spectral clustering, t_o was varied between 0.1 and 0.9 when eigenratio threshold was set at t_elg=5. Alternatively, for spectral clustering, t_elg was varied between 1 and 10 when t_o was set at 0.5. Fraction of assigned alignments (out of 5335 Input) was plotted in panel (C). Fraction of assembled DGs (against 100 Input) was plotted In panel (D). (E) For each simulated DG dataset and clustering parameter combination, the sensitivity and specificity of DG assembly was calculated for each of the top 100 DGs. The sensitivity of DG assembly Is defined as the fraction of remaining alignments in each DG after CRSSANT assembly. The specificity Is defined as the fraction of alignments from the dominant simulated DG. (F) Human U2 snRNA structure model based on previous studies. (G-H) Human HEK and mouse ES PARIS data are clustered using CRSSANT. The DGs were labeled corresponding to the secondary structure models In panel F. Alignments are grouped in IGV using the NG tag. “?” is a new duplex not in the unknown structure models. (I) The duplex SLIld is conserved from human down to yeast based on multiple sequence alignment of 208 seed sequences (Rfam: RF00004, In weblogo format). (J) SLIld model, top strand Is the 5’ arm, while the bottom Is the 3’. Black letters, GUAUGA, Indicate the BPRS masked by SLIld. (K) The alternative SLIM + SLIV structure models.

In crosslink-ligation experiments, crosslinking and ligation efficiencies vary greatly depending on sequence and structure contexts. More importantly, in vivo golden standard structure models do not exist for the vast majority of cellular RNAs since in vitro methods such as cryo-EM, crystallography and NMR only capture a subset of stable conformations under artificial conditions. Therefore, we benchmarked the CRSSANT method on simulated DGs with gap1 alignments (**Supplemental Fig. 4A-B**, Methods and Supplemental Materials). On an artificial chromosome of defined length (e.g. 1000 base pairs), 100 simulated DGs were randomly positioned with defined core length, random extensions on each side of core, random gap length and random numbers of alignments in each DG. Then we clustered the simulated alignments into DGs using the cliques and spectral algorithms and various parameters, including the overlap ratio threshold *t*_*o*_ for both cliques and spectral, and eigenratio threshold *t*_*eig*_ for spectral. *t*_*o*_ was varied between 0.1 to 0.9, while *t*_*eig*_ was varied between 1 and 9. We calculated the fraction of alignments assigned to assembled DGs (**Fig. 4C**), numbers of DGs assembled (**Fig. 4D**), specificity and sensitivity (**Fig. 4E**),

Over 80% of alignments were assembled into DGs with *t*_*o*_ between 0.1 and 0.5 using various simulation settings and both clustering algorithms (**Fig. 4C**). CRSSANT assembly produced between 50 and 200 DGs from the input 100 simulated DGs with *t*_*o*_ between 0.1 and 0.5 (Fig. 4D). As expected, higher *t*_*o*_ (>0.5) reduced assembled alignments, and increased total assembled DGs. Spectral clustering consistently outperforms cliques at the recovery of alignments (**Fig. 4C**), but at the expense of increasing assembled DGs (**Fig. 4D**), leading to unnecessary DG splitting. At *t*_*o*_ above 0.5, performance of both methods dropped, while *t*_*eig*_ did not affect the spectral clustering at all values tested (up to 100, data not shown). Using the optimal settings for the two algorithms (*t*_*o*_=0.5 and *t*_*eig*_=0.5), we examined individual CRSSANT assembled DGs (**Fig. 4E**). More than 80% of the top 100 DGs are consistently assembled with high sensitivity and specificity, i.e., the original simulated alignments are mostly assembled into DGs (sensitivity), and the membership in the assembled DGs are correct (specificity) (all parameter combinations in **Supplemental Fig. 4C-D**). Visual inspection of the assembled DGs confirmed the better performance of the cliques method; the minor reduction of DG numbers compared to simulated input was due to merging of DGs that showed significant overlap at the two arms (**Supplemental Fig. 5A-C**). Even though the spectral method increased recovery of alignments, about 50% of the simulated DGs were split into overlapping smaller DGs (**Supplemental Fig. 5D**). We tested the speed of CRSSANT by varying the simulation and clustering parameters on a standard laptop computer. Increasing alignment numbers in each DG extended running time nearly quadratically since pairwise comparison of overlapping alignments is the bottleneck (**Supplemental Fig. 6A-B**). Consistent with this, increasing genome length while maintaining alignment numbers (effectively reducing alignment density), dramatically lowered running time (**Supplemental Fig. 6C**). The choice of clustering parameters did not affect running time at reasonable overlap thresholds (t_o_ between 0.1 and 0.5, **Supplemental Fig. 6D**). Together, we identified the cliques as the preferred clustering algorithm and showed that t_o_ moderately affect clustering performance.

To validate CRSSANT on cellular RNAs, we analyzed the snRNA U2 and the snoRNA U3 using published PARIS data from human HEK and mouse ES cells (**Lu** et al. 2016) (**Fig. 4F-K, Supplemental Fig. 7**, **Supplemental Table 2** and Supplemental Material). Ungrouped alignments on U2 are densely packed, making it difficult to recognize the structures (**Fig. 4G**). After clustering, DGs have an average dispersion (standard deviations of the left-start, left-end, right-start, and right-end positions) of 5.0 nt for each DG, compared to 44 nt for all alignments on U2, showing that the clustering resulted in tightly packed DGs (**Fig. 4G**). We identified 4 previously known stemloops SLI, SLIIa, SLIII and SLIV (Patel and Steitz 2003; Hilliker et al. 2007; Perriman and Ares 2007). SLIIb and SLIIc were missed due to the lack of psoralen-crosslinkable staggered uridines. In addition, we recovered DGs that suggest new conformations: SLIId and SLIII+SLIV, both of which are conserved between human and mouse (**Fig. 4G-H**). The low-abundance DGs may have come from other less stable conformations (bottom of **Fig. 4G-H**). SLIId is an alternative duplex to SLIIc, masking the branchpoint recognition sequence (BPRS), suggesting a function in regulating U2 recognition of introns. SLIId blocking of BPRS may act as a structural switch to reduce spurious binding and increase splicing fidelity. Analysis of SLIId in U2 revealed a strongly conserved duplex from human to yeast (**Fig. 4I-J**). The near complete overlap of the left arms of SLIII and SLIII+SLIV, and the overlap of the right arms of SLIV and SLIII+SLIV in human and mouse suggest that these conformations are alternative to each other (**Fig. 4G-H**). The left arm of SLIII and right arm of SLIV form a 7bp bulged stem with staggered uridine crosslinking sites, supporting its validity (**Fig. 4K**). Together, the analysis of U2 snRNA validates the CRSSANT clustering strategy, confirming previously known structures, and nominating new conformations that reveal previously unknown mechanisms in splicing regulation.

### Multi-segment alignments provide evidence for complex structures and interactions

Both crosslinking and proximity ligation are inefficient, however, multiple events may occur simultaneously in some RNA regions, leading to reads and alignments with multiple gaps (referred to as gapm, with gaps>=2 or segments>=3, **Fig. 3**). Further analysis of these alignments showed that 3-segment alignments are the majority, accounting for >99% of them, while alignments with more segments were exceedingly rare (**Fig. 5A**). Among 3-segment alignments, ~70-80% of them were mapped within one RNA, while 20-25% of them are mapped to two RNAs simultaneously, indicating RNA-RNA interactions (**Fig. 5B**). A small fraction of them were mapped to 3 different RNAs, suggesting the existence of multi-RNA complexes.

**Figure 5.**
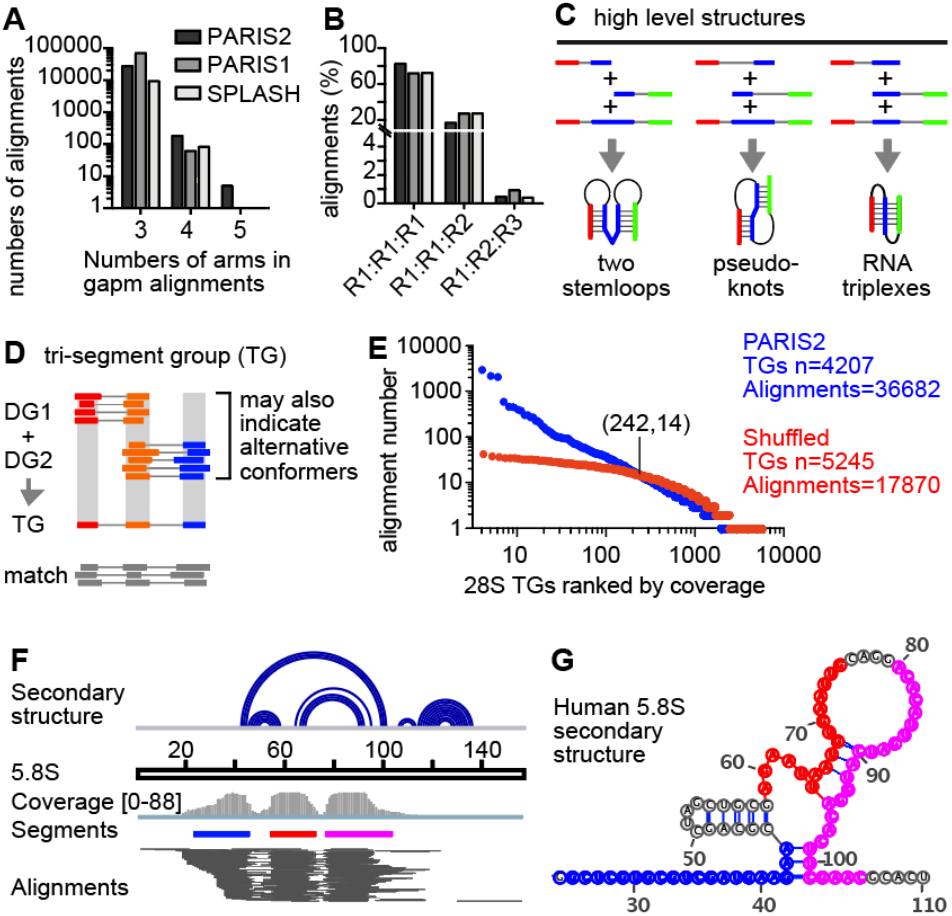
Multi-segment alignments support higher level structures and interactions. (A) Distributions of the numbers of arms/segments in gapm alignments. (B) Numbers of RNAs involved in each gapm alignment. Gapm alignments with 3 arms are shown. R1, R2 and R3 represent 3 different RNAs. (C) Gapm alignments with 3 arms indicate the co-existence of two helical regions. Sequential helices joined by gapm alignments indicate two separate stemloops (left). Interlocked helices joined by gapm alignments indicate pseudoknots (middle). Overlapping helices joined by gapm alignments indicate triplexes. (D) Strategy to cluster gapm alignments, assuming that all TGs should be combinations of DGs. Alignments with more than 2 gaps are ignored for now. The DGs were produced by CRSSANT using gap1.sam and trans.sam alignments. The boundaries for each arm are the medians for the DGs. For the TGs, the merged middle arm is the redefined as boundaries of both DGs. Alignments from gapm.sam are then matched to the TGs so that each arm is overlapped. (E) Gapm alignment number distribution for TGs on the of human 28S rRNA PARIS2 HEK293 gapm alignments were assembled directly on the DGs (blue) or shuffled randomly across the 28S rRNA before assembly (red). (F) gapm alignments mapped to the human 5.8S rRNA. Top track: base pairing secondary structure model in arc format. The 3 segments are color-coded in panels C-D. (G) Mapping the 3 segments to the secondary structure model.

These gapm alignments could indicate several types of structural topology, such as sequential or concentric helices, pseudoknots and even triple helices (**Fig. 5C**, examples in one RNA). For example, we previously showed that interlocking helices suggest pseudoknots, but an alternative explanation is that the two helices could exist in separate RNA molecules (Lu et al. 2016). Alignments connecting the two helices are strong evidence that both helices occur on one RNA, therefore proving the pseudoknot structure. The complex structures could be either intramolecular or intermolecular, indicating complex interactions. Such high-level structures are hard to predict or validate in cells using conventional methods. Focusing on these 3-segment (2-gap) alignments, we developed a method to cluster them into tri-segment groups (TGs, **Fig. 5D**). Given that TGs are combinations of DGs, we first used CRSSANT-assembled DGs to build a list of DG pairs with one overlapping arm. Gapm alignments with 3 segments were then assigned to DG pairs based on overlap with each arm. Three-segment alignments that group together are defined as a TG.

Clustering of TGs from published datasets revealed a large number of complex structures, particularly in the most abundant cellular RNAs, e.g. the rRNAs and snRNAs, and they are consistent with the combinations of DGs (**Fig. 5E** and **Supplemental Fig. 8A-D**). In particular, the top-ranked TGs contain up to 3865 alignments (**Fig. 5E**, blue dots). To test whether TGs correlates with DGs, we shuffled the gapm alignments across the 28S rRNA, and then re-assembled them into TGs. Only 48.7% of the shuffled gapm alignments (17870/36682) can now be assigned. The alignment numbers in each shuffled TG are more uniformly distributed (**Fig. 5E**, red dots), with the maximal coverage at 46, compared to 3865 in the original data. The two distributions crossed at (242,14), where the top 242 TGs contains 75.2% alignments in the original data, but only 27.7% in the shuffled data. These results support the validity of identified TGs. For example, in the 5.8S rRNA, we observed a TG that corresponds to a 3-way junction (**Fig. 5F-G**). Some of the complex structures are supported by more than one TGs. For instance, two concentric helices are supported by 3 different TGs because the RNase cleaved at different locations in the RNA structure before proximity ligation (**Supplemental Fig. 8C**). In addition to intramolecular interactions, we also discovered more complex intermolecular interactions. We previously showed that snoRNAs U8 and U13 form a dynamic network of intermolecular interactions with rRNA precursors during rRNA processing (Zhang et al. 2021). Here we found that gapm alignments connect U8, U13 and the rRNA precursor together, suggesting that these interactions occur simultaneously in cells (**Supplemental Fig. 8E-G**, **Supplemental Tables 4**-**5**). Together, these analyses revealed more complex structures than possible before.

### Identifying alignments with overlapped segments indicating potential RNA homodimers

Base pairing can drive the formation of intramolecular RNA duplexes as well as inter-molecular interactions using the exact same sequences. For example, a stem-loop can also form an alternative conformation of homodimer with nearly identical base pairs (**Fig. 6A**, top and middle). The intermolecular interactions may contain 2 molecules, or even more, forming a daisy-chainlike complex (**Fig. 6A**, bottom). Given the prevalence of RNA stemloops and high concentration of many essential ncRNAs, and the sequestration of mRNAs into RNP granules (Protter and Parker 2016), it is conceivable that such RNA homodimers are widely present in cells. However, homodimers are difficult to detect using conventional methods. Here we found that alignments with overlapping segments enables de novo discovery of such interactions. Normal gapped reads without overlaps between the 2 arms may come from one RNA molecule, or two identical molecules (**Fig. 6B**). Gapped reads with overlaps between them could only have come from a homodimer (**Fig. 6B**). Because of this, such alignments are definitive evidence for homodimers. Such analysis provides an underestimation of the abundance of inter-molecular duplexes since some normal gapped alignments (gap1) may also come from homodimers.

**Figure 6.**
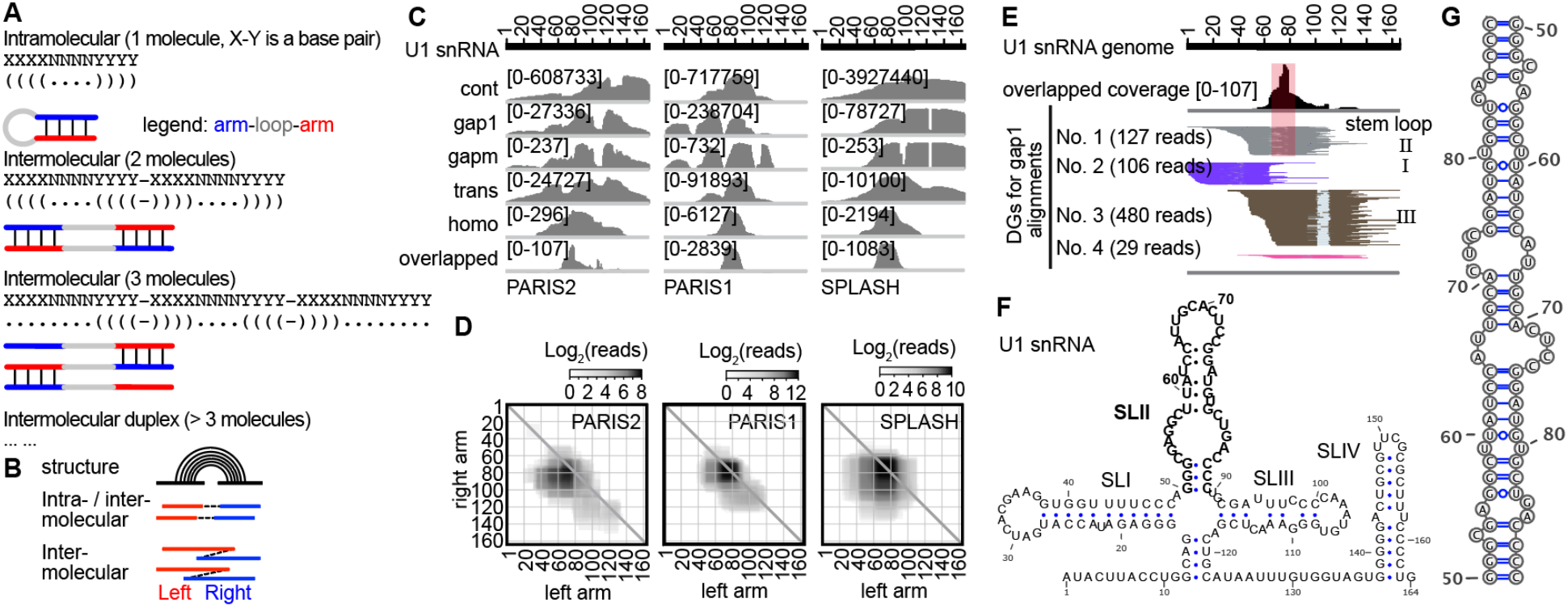
Identification of potential RNA homodimers using homotypic alignments. (A) The same base pairing interactions can mediate intramolecular stemloops (top) and homotypic interactions between 2 (middle) or more (bottom) copies of the same molecule. (B) Diagram showing alignments with gapped or overlapped arms suggesting RNA stemloops or homodimers. (C) Coverage of 5 different types of alignments on U1. The overlapped part of homo alignments was shown individually at the bottom. (D) Heatmap of U1 snRNA homo alignments in 3 datasets. (E) PARIS2 data showing overlapped regions and corresponding local stemloop (SLII). DGs were assembled from 1000 total alignments. (F) Secondary structure of U1 homo interaction, with the SLII in bold letters. (G) Secondary structure model for the SLII homodimer.

To discover RNA homodimers, we analyzed published crosslink-ligation data (summary in **Fig. 3E**). First, we filtered homotypic alignments to remove short 1-2nt overlaps that may come from RNA damages or sequencing errors, and repetitive sequences that may come from enzyme slippage during reverse transcription or PCR. To determine the significance of homodimers, we calculated the ratio of overlapping alignments vs. nonoverlapping ones in the same RNA. Overlapped regions extend to >60nts among various datasets (**Supplemental Fig. 9A**). In general, overlapping alignments are rare, but a few noncoding RNAs have high proportions of overlapping alignments (**Supplemental Table 6**). The most highly enriched RNA is U8, a snoRNA previously shown to be essential for rRNA processing (Peculis and Steitz 1993), mutations in which cause a neurological disease LCC (Labrune et al. 1996; Jenkinson et al. 2016; Iwama et al. 2017). We recently reported this dimer and showed that it is part of a 5 alternative conformations for the U8 snoRNA structure, and this dimer is disrupted by LCC patient mutations [see Fig. 4, S20 and S21 in (Zhang et al. 2021)] (**Supplemental Fig. 9B-D**). In addition to U8, dimers also are likely to form for U1 and U2 snRNAs (**Fig. 6C-D, Supplemental Fig. 9E-I**). In U1, we detected a specific dimerization region in the SLII from 3 different psoralen crosslinking datasets. This specific enrichment compared to broader distributions of other types of alignments further suggests that this homodimer is real, despite the low abundance (**Fig. 6C**). The homo alignments localize to the same sequences as gap1 alignments at the local stemloop (**Fig. 6E**), consistent with them as alternative conformations to each other (**Fig. 6F-G**). Similarly, we detected potential dimerization regions in the SLIII of U2 snRNA, mitochondrial tRNAs, and expansion segments in ribosomal RNAs (**Supplemental Fig. 9J-P**). Overlapping alignments in the mRNAs, however, were not abundant enough to allow the identification of local enrichment sites that indicate dimerization sequences (**Supplemental Table 6**). Homodimerization in other noncoding RNAs may also have been missed due to limited sequencing coverage.

Homodimers have been reported in a variety of RNA viruses. To detect potential homodimers, we analyzed our recently published PARIS2 data on two single stranded RNA virus genomes (Zhang et al. 2021) (**Supplemental Fig. 10**). In both US47 (US/MO/14-18947 and VR1197 (F02-3607 Corn), two strains of EV-D68, we detected local peaks of overlapping alignments. While some of these peaks coincide with local stemloops detected by PARIS2, others were not, suggesting alternative base pairing mechanisms in the interactions (**Supplemental Fig. 10A,D**). The ratio of homotypic alignments over all gapped ones are only ~1% (**Supplemental Table 6**), yet the overlapped regions are rather extended (**Supplemental Fig. 10B-C,E-F**). The top ranked peaks were not conserved between the two viral strains due the rapid evolution of these RNA viruses. Additional dimerization sites may exist that cannot be captured by our method which relies on the identification of local hairpins. Together, these studies demonstrate the ability of our new computational pipeline in the identification of potential RNA homodimers in a variety of contexts.

## Discussion

The recent development of crosslink-ligation methods has dramatically changed the field of in vivo RNA structure studies. Despite the progress in experimental techniques, computational processing of such data remains challenging. Previously developed computational tools have focused on simple cases, i.e., identification of single-gapped alignments and building duplex structures from them (Travis et al. 2014; Sharma et al. 2016; Lu et al. 2018; Zhou et al. 2020). In this study, we performed exhaustive analysis of data from crosslink-ligation experiments, identified limitations of previous computational methods, and designed a set of tools to address several fundamental problems in the analysis pipeline, and to realize the full potential of such experiments.

Specifically, we focused on the mapping, classification, and clustering of sequencing reads. (1). We optimize a set of STAR mapping parameters, together with a new filtering strategy to maximize sensitivity and specificity of aligning short segments. This improvement is particularly beneficial for building higher resolution secondary structure models that require shorter segments. (2). We develop a strategy to exhaustively classify alignments into 8 categories, which are then rearranged to 5 types. The newly developed tools are particularly useful for the analysis of alignments where the two segments can be converted to a single SAM record for visualization in genome browsers (Lu et al. 2016). (3). We develop a network-based method, CRSSANT, for clustering non-continuous alignments to discrete groups that represent the underlying RNA duplexes, for both simple gapped alignments (gap1 and trans), and complex alignments (gapm). We benchmarked each step of the pipeline and demonstrated its applications in various real-world examples. The files output by CRSSANT concisely summarize information that is crucial to the RNA structural biologists and are prepared in file formats commonly used by the structural biology community to facilitate cross-platform analysis. Together this pipeline greatly facilitates the analysis and interpretation of data from a wide variety of crosslink-ligation experiments.

Our systematic analysis of alignment properties, such as the segment length, gap length and gap nucleotide frequencies revealed previously unknown problems that help guide future improvement of crosslink-ligation methods. In particular, we show that the segment length distributions vary greatly across the methods, which has a major impact on the secondary structure modeling. Even with the shortest segments in hiCLIP and PARIS (Sugimoto et al. 2015; Lu et al. 2016), the median segment lengths of ~20 nts far exceed those of the well-studied RNAs such as the ribosome and ribosome (**Fig. 2A**), and it remains challenging to determine the exact base pairs. Future improvements to pinpoint crosslinking sites are necessary for unambiguous modeling. The discovery of psoralen-monoadduct induced uridine deletions, especially in the 1-2nt range, revealed concerns over some of the crosslinking methods. We suggest that these short-gap alignments should be removed before any subsequent analysis.

Even though recent studies have paid attention to alternative conformations in RNA secondary structure modeling from crosslink-ligation data, detailed analysis of individual RNAs is still challenging. In the CRSSANT method, we systematically tested clustering algorithms and parameters on simulated datasets. This benchmarking provides important guidelines for applications on experimental data. As examples, our analysis revealed surprising new conformations, even for well-studied noncoding RNAs, such as U2 and U3. The combination of crosslink data and phylogenetic analysis support the validity of these new conformations, nevertheless, deeper studies are needed to understand their functions and mechanisms of dynamic interconversions.

RNAs in cells are known to form highly sophisticated machines, and our current understanding remains limited to a few well-behaving RNAs and their complexes that can be purified for characterizations. Our exhaustive classification allowed us to discover complex structures and RNA homodimers de novo, further expanding the capabilities of these experimental techniques. In a recent study, we have dramatically improved the crosslinking and overall efficiency of crosslink-ligation experiments (Zhang et al. 2021), however, the low proximity ligation efficiency remains a major bottleneck for crosslink-ligation methods. This problem made it difficult to capture the multi-segment structures and interactions. For example, at 10% ligation efficiency, reads with n segments are less than 1 in 10^n^. Improvement in proximity ligation and the ever-increasing sequencing power should solve this problem to allow the discovery of other complex structures.

The discovery of homodimers is particularly interestingly since it opens new directions for future research. The small fraction of RNAs with overlapping fragments suggest that homodimers based on local palindrome-like sequences are rare. We discovered strong homodimers in the U8 snoRNA, and U1 and U2 snRNAs. These homodimers were detected across different datasets, even though their abundances vary considerably. In the most stable homodimer U8, the overlapping alignments are even more abundant than the intramolecular duplexes in one dataset. Together with our recent in vitro validation (Zhang et al. 2021), we believe that at least a subset of the potential homodimers are real. The discovery of human patient mutations that disrupt the dimers point to functional significance of such interactions (Labrune et al. 1996; Jenkinson et al. 2016; Iwama et al. 2017; Zhang et al. 2021).

We note that in contrast to typical gapped alignments, where shorter segments lead to higher resolution structural modeling, longer segments are needed for efficient detection of overlapping alignments and potential homodimers. In the extreme case of the 5’ end of one copy binding to the 3’ end of another copy of the same RNA, full length RNAs are necessary to detect such dimers. Alternatively, we propose a genetics-based method to detect homodimers, which is not limited by the sequence distance between the two segments (**Supplemental Fig. 10G**). When RNA molecules from two different genetic backgrounds (red and blue lines) exist in the same cell, e.g., during co-infection of two RNA virus strains with sufficient genetic distance between them, or in the F1 generation of a hybrid organism, nucleotide sequence variants allow us to accurately map the fragments to the RNA of origin. When the two fragments are derived from the same genetic origin, the duplex could be either intra- or intermolecular. However, if the two fragments are from two different genetic backgrounds, then the duplex should be intermolecular, i.e., homodimer. Two caveats should be considered in this approach. First, some sequence variations may alter the structures and interactions and lead to artifacts. High enough sequence variation may redefine the homodimer to heterodimer. Second and specifically for RNA viruses, genome recombination may break the linkage of variants, and confound the analysis of intermolecular homodimers.

In this study we focused on the mapping, classification, and clustering of crosslink-ligation data. Subsequent crosslink-guided structure modeling can be achieved using many previously published tools based on free energy minimization and multiple sequence alignments, but it is not a trivial task for several reasons (Eddy 2004; Lu et al. 2016). Even with the shortest fragments available in PARIS, there is ambiguity in determining the exact base pairs. In addition to the problems with the experimental constraints, energy and conservation based computational prediction approaches are still being optimized. As such, manual inspection is still needed for individual RNAs or regions before such models are used guide deeper functional and mechanistic studies. For longer RNAs, it is even more challenging to stitch together all the models derived from individual DGs. Our discovery of the gapm alignments helps resolve certain complex conformations by providing evidence for the coexistence of helices. However, ambiguities also exist in the determination of which fragment base pairs with the other 2 or more fragments in the same alignment (**Fig. 5C**). In a recent study, we reported a new method using computationally enumerated ensembles of RNA conformations, and Bayesian statistics to identify optimal ones that match experimentally determined constraints (Zhou et al. 2020). Further optimization and integration of these various methods has the potential to reveal global conformations for larger RNAs.

In summary, we have developed a new computational pipeline to automate the otherwise laborious tasks of reads mapping, classification, and clustering. We envision that the pipeline will find broad use in the field as crosslink-ligation-based methods are applied to a wide variety of RNA biology problems.

## Methods

### Data access and preprocessing (Fig. 1B, step 1)

All sequencing data used in this study were previously published, and listed as follows. PARIS: *GSE74353* HEK and mES and *GSE149493 HEK* (Lu et al. 2016; Zhang et al. 2021). LIGR: SRR3361013 HEK (Sharma et al. 2016). SPLASH_GMA: SRR3404937, GM12892, polyA. SPLASH_GMT SRR3404942, GM12892 total. SPLASH_ESA: hES (Aw et al. 2016). COMRADES: ZIKV (Ziv et al. 2018). hiCLIP: all data (Sugimoto et al. 2015). Sequencing data from these published crosslink-ligation methods were processed according to the original designs in each study. Briefly, 5’ and 3’ end adapter sequences were removed. Duplicates were removed based on the randomized universal molecular indices.

### Optimized STAR mapping (Fig. 1B, step 2)

The default and optimized formulae for calculating alignment scores are as follows. The major changes are the deletion (deletion), gapopen (gap open), gapext (gap extension) and chimeric junction penalties.

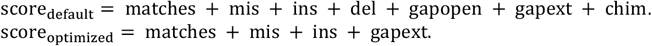

Explanations are as follows. matches: length of matched sequences (+1 for each nt). mis: length of mismatched sequences (−1 for each nt). del: deletion penalty (−2 for deletion opening, −2 for each nt extension). del was disabled by alignIntronMin=1 after optimization, so that all deletions and gaps are considered equal. ins: insertion penalty (−2 for insertion opening and −2 for each nt insertion extention, no change). gapopen: gap open penalty (scoreGapNoncan=−8, scoreGapGCAG=−4, scoreGapATAC=−8). gapopen was disabled after optimization since the gaps are not due to splicing. gapext: gap extension penalty, scoreGenomicLength-Log2scale*log2(genomicLength), changed after optimization. The higher penalty reduces low confidence mapped segments. In chimeric alignments, genomicLength = L1*L2*…*Li, where L is the length of each chimeric segment. chim: penalty for non-chimeric alignment (chimScoreJunctionNonGTAG=−1)

In the final output where all alignments are in the Aligned.out.sam, the counts are defined as follows: primary = unique + chimeric (primary only) + multimapped (excluding ones mapped to too many loci). Here primary alignments can be extracted and counted using samtools view -F 0×900 (Li et al. 2009). An example optimized setting for STAR (Dobin et al. 2013) mapping of non-continuous reads are as follows. --runThreadN 1 (set based on resources) --genomeLoad NoSharedMemory (set based on resources) --outReadsUnmapped Fastx --outFilterMultimapNmax 10 --outFilterScoreMinOverLread 0 --outSAMattributes All -- outSAMtype BAM Unsorted SortedByCoordinate --alignIntronMin 1 --scoreGap 0 --scoreGapNoncan 0 --scoreGapGCAG 0 -- scoreGapATAC 0 --scoreGenomicLengthLog2scale −1 --chimOutType WithinBAM HardClip --chimSegmentMin 5 -- chimJunctionOverhangMin 5 --chimScoreJunctionNonGTAG 0 -- chimScoreDropMax 80 --chimNonchimScoreDropMin 20

### Filtering, classification, and rearrangement of alignments (Fig. 1B, step 3)

The filtering and classification methods are implemented in two scripts gaptypes.py and gapfilter.py. In gaptypes.py, STAR alignments are filtered to remove low-confidence segments, and rearranged and classified to 6 distinct categories (**Fig. 2D-E**, and **Fig. 3**). These six different group alignments were: continous alignments (cont.sam), non-continuous alignments with 1 gap (gap1.sam), non-continuous alignments with multiple gaps (gapm.sam), non-continuous alignments with the 2 arms on different strands or chromosomes (trans.sam), non-continuous alignments with the 2 arms overlapping each other (homo.sam) and non-continuous alignments with complex combinations of indels and gaps (bad.sam). In particular, the fields FLAG, START, SEQ and QUAL, are adjusted after arrangement, while the optional tag fields are left unchanged, except that the ch:A and SA:Z fields that indicate chimeric alignments are removed. See Supplemental Materials for details of the methods.

### Analysis of segment and gap properties (Fig. 1B, step 4)

After mapping and classification, the segment and gap properties are analyzed as a quality control for the data. Specifically, for alignments in sam format, the gap length and segment lengths are summarized and plotted as cumulative densities. At the same time, all sequences in the gap region are summarized to count nucleotide frequencies.

### Removing short and spliced gaps (Fig. 1B, step 5)

In eukaryotes, splicing generate non-continuous reads and alignments and they need to be separated from the ones generated by proximity ligation. We used the annotated splicing junctions to filter the sequencing data as follows. In gap1.sam and gapm.sam, if an alignment only has gaps that are identical to splicing junctions (upper panel), it is removed. If an alignment has at least one gap that is not the same as the splicing junction (lower panel), it is retained. At the same time, all gaps <=2 nts are also removed, since these are highly likely to be artifacts caused by crosslinking induced RNA damages (**Supplemental Fig. 1**).

### Duplex group assembly (Fig. 1B, step 6)

Filtered gap1filter.sam and transfilter.sam alignments are combined as the input. Additional input files are gene annotations in bed format, where only the first 6 fields are needed, and genome files that list sizes of chromosomes. Alignments are assigned to gene pairs based on genome coordinates. If one alignment is contained within one gene (e.g. gene1), then the pair is (gene1, gene1). If the alignment spans gene1 and gene2, then it is mapped to the pair (gene1, gene2). Alignments mapped to each gene pair are processed separately in parallel, to speed up the analysis. Regions with alignments higher than a predefined value are sub-sampled to speed up the processing, and the unused alignments are added back to the assembled DGs. Bedtools is used to produce a genome coverage file from all the 2-segment non-continuous alignments (gap1filter.sam and transfilter.sam). The coverage file is then used to calculate the confidence of each DG. See Supplemental Materials for details about the algorithm.

### Assembly of tri-segment groups (TGs, Fig. 1B, step 7)

Alignments with more than 2 gaps or 3 segments are ignored for now. The DGs were produced by CRSSANT using gap1.sam and trans.sam alignments. The boundaries for each arm are the medians for the DGs. For the TGs, the merged middle arm is the redefined as boundaries of both DGs. Alignments from gapm.sam are then matched to the TGs so that each arm is overlapped.

Specifically, the gapm alignments (mapped to hg38/mm10 genome) were globally annotated using gapm_anno.py scripts. To study the RNA:RNA:RNA structures and inter-molecular interactions from PARIS data, the reads were mapped to selected subsets of RNAs, including snRNA (U1, U2, U4, U6, U5, U11, U12, U4atac and U6atac), highly abundant snoRNA (U3, U8, U13 and U35) and rRNAs. These selected RNA was assembled to one small “chromosome”. After mapping, alignments classification, and short gap filtering, gap1 alignment was used to call RNA:RNA duplex (DG).

The majority of human and mice genome is duplicated sequence, such as repetitive DNA, genes with multiple copies. This makes unambiguous identification of RNA:RNA:RNA interactions very difficult on a genomic scale. To identify the RNA-RNA-RNA structures and interactions from PARIS data, the reads were mapped to selected subsets of RNAs, including snRNA (U1, U2, U4, U6, U5, U11, U12, U4atac and U6atac)., highly abundant snoRNA (U3, U8, U13 and U35) and rRNAs. These selected RNA was assembled to one small “chromosome”. After mapping using STAR program, alignments was classified into 6 groups. Filtered gap1 alignments (gap length > 2nt) was used to call DGs. The assembled gap1 DGs were further used to cluster gapm alignments. The curated DGs were used for TGs assembly for gapm alignments. U8:U13:28S inter-molecular interaction were analyzed using PARIS1 mES data (GSM1917758, GSM1917759 and GSM1917760) (Zhang et al. 2021).

### RNA homodimer (homo.sam) analysis

Homo alignments (homo.sam) with less than 2nt overlapping between two arms were filtered out to avoid potential artifacts. The distance between two arms was calculated: overlap = min(arm1_end, arm2_end) − max(arm1_start, arm2_start). To understand the relationship between RNA homodimers and RNA stem loop structures. RNA stem loops were identified using local gap1 alignments. The length of two arms should be larger than 15nt and the loop length (gap length) should be less than 20nt.

### Bowtie mapping of alignments and subsequent processing

To compare the STAR and Bowtie2 mapping protocols, we designed the following general pipeline to map reads using Bowtie2 and process the alignments. First the reads were mapped using two sets of published parameters (Travis et al. 2014; Sharma et al. 2016). The alignments were converted to chimeric format using bowtie2chim.py, a custom script to rearrange chimeric alignments. The rearranged alignments were filtered using gapfilterbt2.py to remove splicing events, and unique alignments with deletion (D in CIGAR) >2 are counted. Similar to the STAR mapped data, the gaptypes.py script was used here to classify the alignments.

Brief description of the bowtie2 parameters. D: seed extension attempts, R: reseeding attempts, N: max mismatches, L: seed length, k: max number of valid alignments to search, i: score-min: ma: match bonus, np: N penalty, mp: mismatch penalty, rdg: affine read gap penalty, rfg: affine reference gap penalty. For end-to-end mode, the minimum should be −0.6-0.6*L, where L is the length of the read. For local mode, the minimum should be 20+8*ln(L). The commonly used setup in bowtie2:

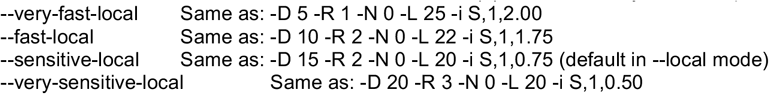

The primary assembly of hg38, and combinations of the Rfam and annotated mRNAs were used to build the bowtie2 indices (Lu et al. 2016). The Bowtie2 parameters from the hyb package are as follows (Travis et al. 2014): bowtie2 -p 20 -D 20 -R 3 -N 0 -L 16 -k 20 --local -i S,1,0.50 --score-min L,18,0 --ma 1 --np 0 --mp 2,2 --rdg 5,1 --rfg 5,1 -x hg38pri -U xxx.fastq -S xxx.sam. The Bowtie2 parameters from the Aligater package are as follows (Sharma et al. 2016). bowtie2 -p 20 -k 50 -R 3 -N 0 -L 16 -i S,1,0.50 --local -x hg38pri -U xxx.fastq -S xxx.sam. Default for the other parameters are as follows: -D 15-score-min G,20,8 --ma 2 --np 1 --mp 6,2 -- rdg 5,3 --rfg 5,3. The rdg and rfg setting in the default is much stronger.

Bowtie2 does not produce supplementary alignments like STAR. It has one primary and multiple secondary alignments ("FLAG 256"). In the unsorted sam output, the first is always the primary alignment. We developed the following strategy to combine the primary and secondary alignments (bowtie2chim.py). If any of the secondary alignment can combine with the primary, with at most overlap (eg. 2nt) in the query, then we consider the pair as a chimera, and modify the two alignments with tags ‘ch:A:1\tSA:Z:A’. If multiple loci for secondary alignments can be matched to the primary, keep one for simplicity. Only reads without linkers are considered to be comparable to a typical STAR run. The linkers can be easily removed before the STAR mapping step.

### Simulation of DGs to benchmark classification of gapped alignments

First, on an artificial chromosome of chr1:0-1000, made of nucleotides “N” in one gene GENE1: 0-1000, pairs of core intervals of a specified length (corelen, e.g. 5, 10 or 15nts) are selected, in the range of chr1:100-900. The first 10kb of hg38 chr1 happens to be a stretch of “N”, so the results can be viewed on hg38. The two intervals of each pair are at least coregap away from each other (e.g. coregap=50), and within a specific distance (e.g. corewithin=1000, for chr1:0-1000). Each side of the two cores are extended by a random length (e.g. in the range [5,15]) to make one alignment. Each pair of core intervals are expanded to a set number of alignments that make up one DG, and the number of alignments in each DG is randomly set in a specific range, e.g. DGlower=10, DGupper=100. A set number of DGs are generated (e.g. 100), and overlap between DG cores are allowed for at most one arm, but not both. Pseudo random numbers were generated with seeds to ensure reproducibility. This script, dgsim.py generates a simple set of alignments in DGs to test crssant.py.

### RNA Secondary structure modeling and visualization

In general, base pairing was predicated using ViennaRNA Package (v 2.1.9)(Lorenz et al. 2011). DGs and TGs alignments were visulized by Itergrative Genomics Viewer (IGV, v2.8.13)(Robinson et al. 2011). The curated seed alignments were turned into a weblogo (https://weblogo.berkeley.edu/logo.cgi). Each arm of gapm and homo alignments were mapped to human 28S rRNA cryo-EM structure (ID: 4V6X), U1 snRNP cryo-EM structure (ID: 3CW1) and U2/U5/U6 snRNP cryo-EM structure (7ABI).

## Software availability

Source code for the software developed in this study and is available from GitHub (https://github.com/zhipenglu/CRSSANT).

## Acknowledgements

We thank members of the Lu lab and C.A. Weidmann for discussion. The Lu lab is supported by startup funds from the University of Southern California, the NHGRI Pathway to Independence Award (R00HG009662), NIGMS (R35GM143068), USC Research Center for Liver Disease (P30DK48522), Illumina and USC Keck Genomics Platform (KGP) Core Lab Partnership Program, the Norris Comprehensive Cancer Center (P30CA014089) and USC Center for Advanced Research Computing.

## Author contributions

M.Z., I.T.F.-H., T.W. J.Y.Z and Z.L. conceived and designed the project. M.Z., I.F.-H. and Z.L. wrote the software with input from T.W. and J.Y.Z. M. Z. K.L. and Z.L. performed the data analysis. M.Z., I.F.-H. and Z.L. wrote the manuscript with input from all other authors. Z.L. supervised the project.

## SUPPLEMENTAL MATERIAL

### 1. Annotation and optimization of STAR parameters related to non-continuous alignments

Given the challenges in the analysis of gapped alignments (**Supplemental Table 1**), here we describe the choice and settings for all the relevant STAR parameters, out of more than 100 (version 2.7.1a, May 14, 2019) (Dobin 2019). A subset of these parameters has been discussed (supplemental methods in (Lu et al. 2016; Lu et al. 2018)). Some of the following discussions were from Dr. Dobin in online forums. The most recent SAM/BAM format was used (Sequence Alignment/Map Format Specificiation 2019-05-14, Sequence Alignment/Map Optional Fields Specification 2019-01-30).

**Supplemental Table 1.**
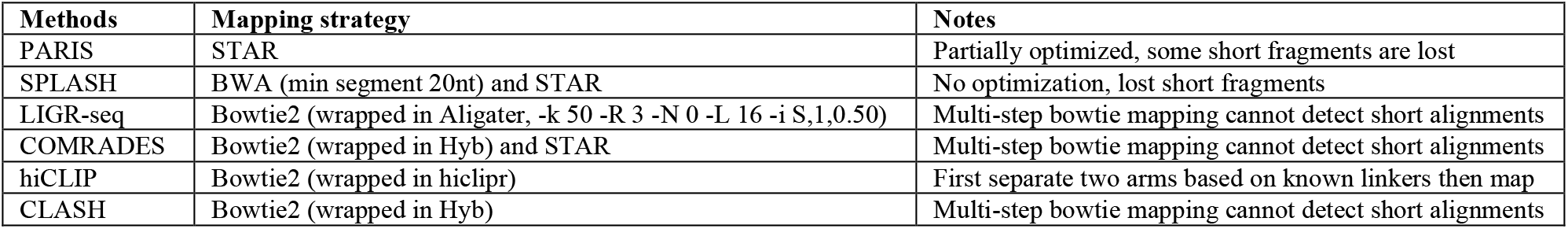
Mapping strategies of published crosslink-ligation studies.

All parameters that are either related to non-continuous alignments or recommended for best performance or are selected from section 14 of the STAR manual and discussed below. Values other than defaults are suggested for optimal use. Additional discussions about genome references are presented in the following sections. The most critical parameters that affect gapped and chimeric alignments are in sections 14.17 Scoring, 14.18 Alignments and Seeding (especially for small genomes), and 14.21 Chimeric Alignments.

For parts of a read aligned within a genome window, the following parameters critically affect the stitching and scoring process and are set specifically for the analysis of data from PARIS and related methods: out*, scoreGap*, alignIntronMin and chim*. The two parameters chimSegmentMin and chimSJoverhangMin affect the identification of gapped reads that span multiple genome windows. They are similar, but the chimSJoverhangMin covers the situation where a splice junction is near the chimeric junction (Haas et al. 2017).

### 14.3 Run Parameters

These are useful for customization of the running environment.

--runMode genomeGenerate for making new indices, see section 14.5 for more details on generating indices for nontypical genomes. alignReads for mapping. liftOver for lifting GTF files among different assemblies.

--runThreadN, use more threads to speed up mapping when running the job in a large computer cluster.

### 14.4 Genome Parameters

Standard ones that are not critical for the analysis are not discussed here.

--genomeLoad LoadAndKeep for fast loading and mapping of reads to large genomes like human. Use other values accordingly. On the fly junction insertion and 2-pass mapping cannot be used with shared memory genome.

--genomeSAindexNbases, define based on genome size, see section 14.5 for its use in generating genome indices.

--genomeChrBinNbits, defined based on total reference size and number of “chromosomes” in reference. See section 14.5 for its use in generating genome indices.

### 14.5 Genome Indexing Parameters, only used when generating genome indices

STAR will not run without the correct parameters in this category. See appendix A2 and A2 on details of mapping to small genomes

--genomeChrBinNbits default 18. Set as log2(chrBin), should be adjusted for a genome with a large number of contigs, such as mapping to the transcriptome, using the following formula: min(18,log2[max(GenomeLength/NumberOfReferences,ReadLength)])

--genomeSAindexNbases default 14. For small genomes, this parameter must to be scaled down, with a typical value of min(14, log2(GenomeLength)/2 - 1).

### 14.6 Splice Junctions Database

--sjdbFileChrStartEnd, This is not the recommended method for removing spliced reads as many gapped reads that span splicing junctions are useful for structure analysis. Including this reference information may be useful for other purposes.

--sjdbGTFfile, allows conversion of genomic coordinates to each transcript to simplify subsequent analysis on the transcript level. For example, converting mapped reads to each transcript allows facile analysis of percentage of gapped reads. This method may also obliviate the need to remap the reads to the transcriptome separately. Mapping reads using genome indices generated with GTF annotation files is faster than including the annotations on the fly. However, using the annotations on the fly is more flexible since multiple annotations can be used separately.

### 14.9 Read Parameters

--readFilesType, default Fastx (fasta and fastq). “SAM SE” and “SAM PE” can be used to input SAM file for remapping reads, without converting reads back to fasta or fastq.

--readFilesCommand, commandline to execute for each of the input files, can be used to decompress files. This command is convenient when input files are zipped to save space.

--clip*, these parameters may be used in place of other adapter trimming software. Use according to need. Removing adapters before running STAR is recommended.

### 14.10 Limits

--limitOutSJcollapsed default 1000000 (1M). We usually need to set a bigger value for larger dataset with higher amounts of non-continuous reads, especially for datasets where the fractions of non-continuous alignments are high. May trigger segmentationfault. See section 3.3 for more details about allocating more shared memory.

--limitIObufferSize default 150000000 (150M). This parameter should be adjusted proportionally to the limitOutSJcollapsed (see discussion here: https://groups.google.com/forum/#!topic/rna-star/FyCj6ZB_tF8). For example, the limitIObufferSize parameter can be set at 150 times of limitOutSJcollapsed. However, this number was deduced from Dobin’s answers cited above, we are not sure how this should be set. The parameters may need to be tested when working with large datasets with high percentage of gapped reads. For example, if we have 100M junctions, then would need 15G IO buffer per thread.

### 14.11 Output: general

--outFileNamePrefix, need to set a prefix, otherwise all output will have the same names and overwrite previous runs.

--outSAMattributes All, needed for duplex group (DG) assembly later, especially the SA:Z tag, which indicates chimeric alignments.

--outReadsUnmapped, default None. Set as Fastx so that the unmapped reads can be examined for troubleshooting.

### 14.12 Output SAM and BAM

--outSAMtype default SAM. Setting this parameter as BAM saves time in later step of SAM to BAM conversion which is quite slow. The “Unsorted” option preserves the order of reads, but the SortedByCoordinate option further saves time.

### 14.15 Output filtering

--outFilterMultimapNmax default 10, needs to change according to genome index used. For example, 1 is recommended for a minigenome without repetitive sequences. Primary alignments can be extracted later using samtools view -F 0×900.

--outFilterScoreMinOverLread default 0.66. Set this to a smaller value, for example, 0, when imposing high extension penalty to discourage long gaps with short matches. See section 14.17 for gap scores.

--outFilterMatchNminOverLread default 0.66, need a lower number, e.g. 0.5, similar as above if there are long unmapped parts in reads. The lower thresholds for these two parameters will capture more gapped alignments, especially in cases where the softclip portion is big.

### 14.16 Output filtering: splice junctions

--outSJfilterOverhangMin default 30 12 12 12 for noncanonical, GT/AG and CT/AC motifs, GC/AG and CT/GC motifs, AT/AC and GT/AT motifs. These parameters only affect the output in the SJ.out.tab file, but not the Aligned.out.sam file. They may be useful when subsequent analysis uses the SJ.out.tab file. Other parameters in this category are used in a similar manner (see discussion here: https://groups.google.com/forum/#!msg/rna-star/J6qH9JCysZw/VmgoGKSE9qQJ). Since the SJ.out.tab file is not used in PARIS analysis, changing these parameters is not necessary at the moment,

### 14.17 Scoring

These parameters are related to outFilterScoreMinOverLread and should be set together (see section 14.15). alignSJoverhangMin determines the minimal overhang size, which is 5 by default. It is related to the scoring but does not need to be changed (see section 14.18). Together, the new settings for scoreGap* and scoreGenomicLengthLog2scale decreases gap opening penalty while increasing the gap extension penalty.

--scoreGap, default 0. Setting this to a large negative value to discourage gap opening when mapping to small genomes is not a good approach as it affects both short and long gaps the same way. Deletions are not considered when setting alignIntronMin=1

--scoreGapNoncan, default −8, set to 0

--scoreGapGCAG, default −4, set to 0

--scoreGapATAC, default −8, set to 0. These three parameters are set to the same value for consistency, since gapped reads from proximity ligation are not the result of splicing.

--scoreGenomicLengthLog2scale, default −0.25. The product of scoreGenomicLengthLog2scale*log2(genomicLength) is added to the penalty. genomicLength should not be confused with genome length. The maximal genomicLength is limited by window size (2^winBinNbits)*winAnchorDistNbins, or 589824 by default (see section 14.20 for details). Therefore, the maximal penalty is scoreGenomicLengthLog2scale *log2(589824), log2(589824) = 19.17. For a ~20kb transcript, such as the human XIST RNA (19287nt), log2(20000) = 14.29. In order to reduce gap openings that lead to small matched fragments, this parameter can be set to a maximal value without discouraging too many long-gap alignments. Setting this parameter to 1 will enable mapping of reads with long gaps with a smaller match of 20 and 15nt. These are reasonable lengths considering the increased chance of errors for a fixed fragment on a larger reference lengths. With the default setting of −0.25, the penalty will be only −4.79 and −3.57, which are too small to discourage gap opening. Note, * is used both as multiplication and wildcards in this document.

--scoreDel* are disabled by setting alignIntronMin to 1 (see section 14.18).

--scoreIns* are set at default values −2.

### 14.18 Alignments and Seeding

The seeding parameters can be altered to affect the alignment behavior, but are too complex to use effectively.

--alignIntronMin set to 1, this would shift all deletions (CIGAR: D) to gaps (CIGAR: N, like introns) and level the playing field in penalty. Gap penalty is calculated as sum(scoreGap*, scoreGenomicLengthLog2scale*log2(genomicLength)). Possible CIGAR operations include “MINSH” (alignment match, insertion, gap, softclip and hardclip), but not “DP=X” (deletion, padding, sequence match and mismatch). Deletion penalty is calculated as sum(scoreDel*), which is 0 when deletions are shifted to gaps. See section 14.17 for details.

--alignIntronMax, default is (2^winBinNbits)*winAnchorDistNbins, or 589824, same as the window size, This parameter is smaller than many genes, and we may need to set it to a larger number in order to capture long-range structures in PARIS data. Check gene size distribution: https://www.biostars.org/p/43283/. For example, the human dystrophin gene DMD covers 2.3 megabases. Haas et al. suggests 200000 for this parameter (Haas et al. 2017).

--alignMatesGapMax, similar as above, these two determine whether local stitching occurs. Mapped fragments that span beyond this range will be considered as chimeric.

--alignSJoverhangMin default 5, should be enough for PARIS and related methods. A larger number will eliminate short overhangs when mapping to small genomes, however, this equally affects short and long gaps, and lead to loss of reliable gapped alignments with short gaps.

--alignSplicedMateMapLminOverLmate, default 0.66.

### 14.20 Windows, Anchors, Binning

These parameters are relevant to mapping gapped reads, but they are statically set. In the analysis of non-continuous alignments, we need to dynamically define the reliable window for gaps, so changing these parameters do not help.

--winBinNbits, default 16. Int >0: = log2(winBin). This parameter determines the bin size for windows and clustering.

--winAnchorDistNbins, default 9. The default window is then calculated as (2^winBinNbits)*winAnchorDistNbins = 589824.

### 14.21 Chimeric Alignments

--chimOutType, two options: WithinBAM and SeparateSAMold. The latter option was the default in earlier versions but will be deprecated in the future. Setting WithinBAM HardClip has several benefits: primary alignments can be calculated easily with samtools view -F 0×900; penalty-based filtering can be performed together and with consistency. Two options are available: HardClip and SoftClip. The HardClip only applies to one of the pair of chimeric alignments, which is useful for marking chimeric alignments.

--chimScoreSeparation (=10 by default) determines the minimum difference (separation) between the best chimeric score and the next one. If the difference is less than this value, the chimera is considered multimapping and is not output. This parameter applies for the old chimeric detection (with default --chimMultimapNmax 0). The new chimeric detection with --chimMultimapNmax >=1 can detect multimapping chimeras uses -- chimMultimapScoreRange (=1 by default) instead.

--chimSegmentMin default 0, recommend 5. This parameter is critical for three reasons. First, setting a value other than the default 0 switches on the chimeric mapping procedure. Second, the value determines how many chimeric reads are reported, and the confidence. Third, smaller chimeric alignments can be filtered out later based on distance to the first larger one. The filtering parameters will be similar to the gap extension penalty to make it consistent.

--chimScoreJunctionNonGTAG (default is −1) int: penalty for a non-GT/AG chimeric junction. Should be set to 0 since the chimeric reads are not due to splicing.

--chimMultimapNmax, the maximum number of chimeric hits

--chimJunctionOverhangMin, recommend 5, similar to the chimSegmentMin parameter, but deals with splice junctions at the same time.

--chimNonchimScoreDropMin default 20.

The following are explanations by Dobin (https://groups.google.com/forum/#!topic/rna-star/CIX-PKCXdCo): “The ChimScore is calculated as the sum of the scores of chimeric fragments. Then the chimeric alignments are output if (ChimScore >= --chimScoreMin) AND (ReadLength-ChimScore > --chimScoreDropMax) AND (ReadLength-BestNormalAlignment > chimNonchimScoreDropMin), where ReadLength is the sum of mates’ lengths for PE reads, BestNormalAlignment is the best score for the normal (non-chimeric) alignment for this read.” However, I would like to point out that the second “>” is likely a typo and should be “<=”.

ChimScore is calculated as follows, here ignoring the mismatches, gaps, indels and chimScoreJunctionNonGTAG, only considering genomicLength penalty: L + S + scoreGenomicLengthLog2scale *log2(L*S), where L and S represent the lengths of the long and short mapped chimeric fragments. This penalty is obviously larger than for a normal full length nongapped alignment: L + S + scoreGenomicLengthLog2scale *log2(L+S), because L*S>L+S for normal match lengths (for example 5-50nt). Therefore the criteria described above can be represented as follows. First of all it is unlikely that an alignment will have ChimScore < 0, unless in rare cases there is a big penalty from additional gaps. So only the second and third ones are considered here:

ReadLength-ChimScore = -scoreGenomicLengthLog2scale *log2(L*S) <= chimScoreDropMax (default 20). Here we already set the chimScoreJunctionNonGTAG value to 0. It follows that L*S <= 2^(chimScoreDropMax/(-scoreGenomicLengthLog2scale)). After changing the value of scoreGenomicLengthLog2scale from −0.25 to −1, I also need to increase the chimScoreDropMax to a much larger value. Setting a value of 80 should cancel the effect of altering scoreGenomicLengthLog2scale, but this is way more than enough, because it produces a score of L*S <= 2^80. It will never reach that for PARIS and related data.

ReadLength-BestNormalAlignment = L+S-(L+scoreGenomicLengthLog2scale*log2(L)) >= chimNonchimScoreDropMin (default 20). Given the increased absolute value for scoreGenomicLengthLog2scale (negative), this criterion is easier to fulfill. Therefore increasing chimNonchimScoreDropMin will increase stringency. However, so far we do not have any problem.

### 14.22 Quantification of annotations (output in transcript coordinates)

--quantMode default-. The TranscriptomeSAM mode should be used together with the inclusion of the transcriptome reference. This mode is useful for subsequent analysis of RNA structures on the transcript level. However, this option cannot handle repetitive sequences for many important noncoding RNA, such as snRNAs and rRNAs. Examining intermolecular interactions requires remapping to the transcriptome annotation because many noncoding RNAs have multiple copies in the genome, which reduces detection sensitivity.

--quantTranscriptomeBan, default IndelSoftclipSingleend. Surprisingly, setting the quantMode parameter does not output gapped reads in the transcriptome coordinates. This is probably a bug in the current STAR version. So it is not quite useful for PARIS and related methods.

A summary of the optimized STAR parameters is listed in **Supplemental Table 2**.

**Supplemental Table 2.**
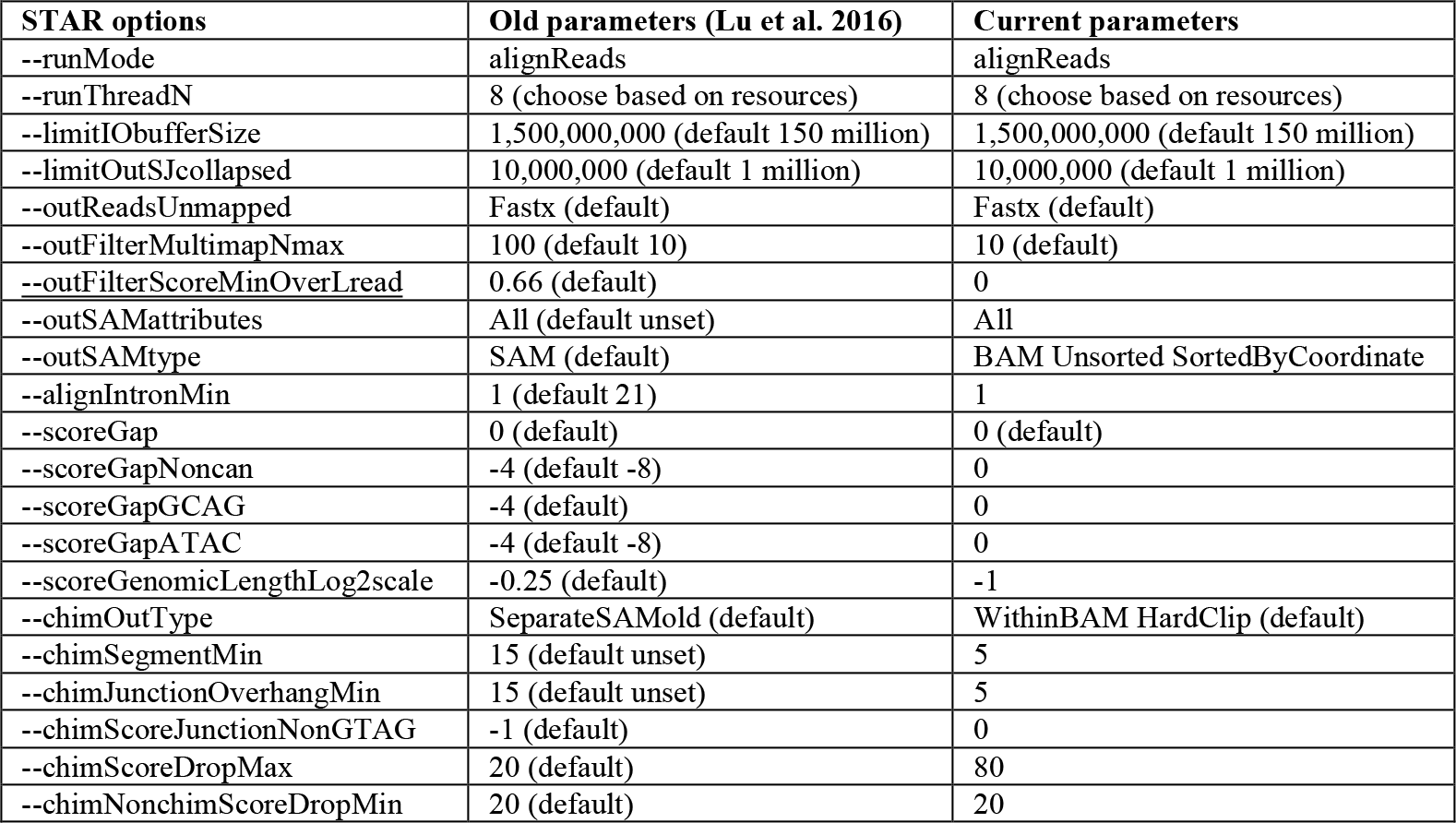
Comparison of STAR settings.

Summary of the optimized STAR parameters for penalty calculation

1. mismatches: −1
2. insertion (I) open: −2
3. insertion (I) ext: −2
4. deletion (D) disabled
5. gap (N) opening: 0
6. map length: xlog2(L)
7. chimeric junction: 0

## 2. Applying STAR to genomes that are either small or contain large number of chromosomes

STAR was designed and optimized primarily for mammalian genomes (Dobin et al. 2013), genomes that are large and contain a small number of chromosomes or contigs (e.g. 3G bases and 194 contigs for human genome primary assembly of hg38). In reality, it is important and useful to map reads to small genomes or large numbers of contigs. In the case of PARIS and related methods, we often align reads to a minigenome containing a single gene, or a small number of genes, or the entire transcriptome annotation that contains tens of thousands of genes, each as a “chromosome”. Relevant parameters described above will be redundantly described here to facilitate the understanding.

1. In the annotations of STAR parameters, we pointed out several approaches for reducing short overhangs in mapping to small genomes. Increasing scoreGap (gap opening, e.g. from 0 to −30) imposes a penalty that affects all alignments equally, and as a result, many good alignments are lost due to this choice (short gaps and short overhangs, e.g. 20M10N10M). Similarly, increasing alignSJoverhangMin, from 5 to 10 or 15, removed all short overhangs. The genomicLength based penalty is more reasonable, since the penalty is log scaled roughly to gap length.
2. Mapping reads to small genomes is much slower than mapping to the whole genome. Dobin explained this problem in a reply to questions, https://groups.google.com/forum/#!topic/rna-star/cLpf7BuDnGY. “The problem is not in smallness of the genome per se, but rather in the incompleteness of the genome. Since majority of the reads will not map to the rRNA reference, STAR would be trying hard to place them, which slows down the mapping speed. I think including the rRNA sequences with the human genome is the best option. You are right, it will increase the multi-mapping for some reads, however, you can deal with such alignments in the postprocessing. Most of rRNA alignments will be multi-mappers anyway. Another option is to map to the standard genome first, and then re-map the unmapped reads (output with --outReadsUnmapped Fastx) to the rRNA reference.” Based on this principle, it is recommended to map reads to a collection of transcripts. We use the RfamhumanrnaMrna and RfammousernaMrna collections. The RfamhumanrnaMrna collection contains 26469 contigs (genes/RNAs), total 83624264nt, while the RfammousernaMrna contains 26967 contigs, total 73085539nt.

## 3. Choice of genome references and annotations for STAR mapping

Genome indices are regenerated from primary assembly fasta files. Primary assemblies recommended for STAR mapping since alternative patches cause multiple mapping. Currently we use hg38 reference fasta from IGV after removing the alternative contigs since they are very similar to existing sequences in primary assembly (see this for details: https://software.broadinstitute.org/gatk/documentation/article?id=11010). This reference contains 194 contigs and is easy to use in IGV and associated annotations. To download the fasta from the following route: Genomes/Load Genome From Server, choose the option: Download Sequence. To remove the alternative patches, use script hg38_igvprimary.py. The Ensembl reference released on 2019-05-24 include assembled chromosomes, unlocalized and unplaced sequences (Homo_sapiens.GRCh38.dna.primary_assembly.fa.gz), but chromosome names are in a different format.

All the reference files are located here: https://s3.amazonaws.com/igv.org.genomes/. The source of this information is here: https://github.com/igvteam/igv/blob/master/genomes/genomes.txt. The additional URL key for hg38 is hg38/refGene.sorted.txt.gz. This file contains all the variants visible in IGV, including 27845 genes (oe) and 71644 transcripts. To convert this file to GTF format for STAR mapping, use the following script. cut -f 2-hg38_refGene.sorted.txt |./genePredToGtf file stdin hg38_refGene.sorted.gtf. The gene count is much lower than the Ensembl annotation of 52506 genes (coding + noncoding + pseudogenes) and 226950 transcripts in the primary assembly (https://uswest.ensembl.org/Homo_sapiens/Info/Annotation, Homo_sapiens.GRCh38.97.chr_patch_hapl_scaff.gtf). This annotation is way too complicated than necessary. For example, it includes 18 isoforms of ACTB, 16 isoforms of ACTG1, 17 isoforms of MALAT1, 30 isoforms of XIST, way more than normal. In total 247909 transcripts are annotated. See this discussion for details: http://www.acgt.me/blog/2015/4/29/how-can-you-choose-a-single-isoform-to-best-represent-a-gene.

The mouse genome annotation at Ensembl contains 52332 genes (coding + noncoding + pseudogenes) and 142333 transcripts in the primary assembly (https://uswest.ensembl.org/Mus_musculus/Info/Annotation). In the IGV RefGene on the contrary has much fewer genes and transcripts, 24982 (20787 NM and 5298 NR) and 42574 (36145 NM and 6429 NR) respectively.

## 4. Filtering, classification, and rearrangement of alignments

In practice, normally arranged (types 1-3) and chimeric (types 4-7) alignments are output to two files (Aligned.out.sam and Chimeric.out.sam) in earlier versions of STAR, with some chimeric alignments in both files, causing confusion. In more recent versions of STAR, all types of alignments can be directed to the Aligned.out.sam file using the chimOutType WithinBAM HardClip option, simplifying the analysis. Due to limited ligation efficiency, continuous alignments represent the majority of the sequencing output, and are removed from further analysis. All the rest (types 2-8) are defined as non-continuous and processed as follows.

### 4.1.

All non-continuous alignments where at least one segment is shorter than a predefined value are first filtered out (gaptypes.py). If the length of the short segment is longer than abs(log2(genomicLength)*scoreGenomicLengthLog2scale), the short segment is valid. This criterion is the same as the gap extension penalty during STAR mapping. In other words, if a short segment is close to a long segment in the same alignment, it is likely to be unique and authentic. Otherwise, the short segment is tested against a database of confident non-continuous alignments in the following step (**Fig. 2D-E**).

### 4.2.

We build a database of confidant connections based on non-continuous alignments (**Figure 2E**). All segments that are equal to or longer than a predefined value (e.g. 15nt), are used to build the connection database. For each segment interval, positions that end with 0 or 5 are selected for building an all-vs.-all connection database. This sparse grid is used to reduce the database size, by ~25-fold, speeds up the search by 25-fold, while maintaining the resolution of the database, because all alignment segments are longer than 5nt are guaranteed to have at least one query chance. The format of the connections is a tuple: (RNAMEi, STRANDi, POSi, RNAMEj, STRANDj, POSj). In practice, we take all POS in the range [START-4, END+4], inclusive, that end with 0 or 5. For the search stage, only query POS in the range [START, END], inclusive, that end with 0 or 5. For example, in segment, chr1, ‘+’ 97-114, positions 95, 100, 105, 110 and 115, are selected to build the database. Here is an example connection: (‘chr1’, ‘+’, 100, ‘chr2’, ‘-‘, 205’), which means that position 100 on the plus strand of chromosome 1 is connected to position 205 on the minus strand of chromosome 2. The database is implemented in a hash table for rapid query. If non-continuous alignments have short segments that match the database, they are considered valid, otherwise, the short segments are trimmed. For alignments that have multiple segments, only the shortest one is tested this way. This database-dependent filtering method is like mapping reads to a genome reference that contains splice junction information, where even a short overhang can be accurately captured. For example, a spliced alignment 20M10000N5M is valid if it precisely aligns to a splicing junction. Like the de novo curation of a splicing database, sequencing depth affects the size of the connection database, therefore a larger one can be generated from a better dataset and used for filtering smaller sequencing datasets.

### 4.3.

Converting type 3, distant collinear chimera, to type 2 (**Fig. 3B**). In STAR and other short read mappers, collinearly aligned segments are considered chimeric. The distinction between types 2 and 3 is determined by the window size, an arbitrary parameter in mapping. To avoid confusion, we converted type 3 to type 2, the distance between the two segments is calculated and then used to connect to two separate segments. This conversion allows the two SAM records to be combined and represented in one line.

### 4.4.

Converting type 4, backward chimera, to type 2 (**Fig. 3C**). Chimeric alignments where all segments are on the same strand and the same chromosome (types 3 and 4) are rearranged so that they can be represented by a single record and CIGAR string. For backward chimera, the two segments are switched so that they are in forward arrangement just like type 2 (**Fig. 3A**). After rearrangement, the two segments of one alignment can be visualized properly in genome browsers, like IGV. In theory, there should be roughly similar number of reads in backward chimeric (type 4) and normal gapped categories (types 2-3). Therefore, rearrangement of the backward chimeric to normal gapped is important for highly sensitive detection and visualization of RNA duplexes. In practice, the backward chimeric (type 4) are less than the normal gapped (type 2), because mapping is biased towards normal gapped.

### 4.5.

Converting type 5, overlapping chimera, to a single CIGAR representation. Chimeric alignments where two arms overlap cannot be represented by a single record. This limitation makes it difficult to visualize them on genome browsers. Here we introduce a new approach for this type of chimeric alignments. In the SAM format, ‘I’ (insertion) consumes query but not reference, while ‘D’ (deletion) consumes reference but not query, therefore 2I1D will consume 2nt query but only 1nt reference, corresponding a 1nt overlap in the middle (**Fig. 3D**). The example new CIGAR 20M2I1D2I1D2I1D2I1D8M can also be represented as 20M8I4D18M, but the former is better visualized on IGV. These overlapping chimeras could be generated because of RNA homodimers and we noticed there are many potential homotypic interactions in PARIS and similar methods, especially when the arms are longer.

## 5. CRSSANT method details (clustering gap1 and trans alignments to DGs and NGs)

### 5.1 CRSSANT data pre-processing

The mapped reads/alignments accepted as input to the CRSSANT pipeline are in the SAM file format (Li et al. 2009). Currently only alignments with two arms are used, i.e. gap1.sam and trans.sam output from the previous step of classification, with splicing alignments removed. These alignments represent the majority of non-continuous alignments. To cluster these reads/alignments, they must be first converted into 4-tuples of alignment arm start and stop indices. This is done using the alignment start position and its CIGAR alignment string, which encodes the transformations performed on the read during alignment. During the ligation step in the PARIS method, ligation may occur in two different ways. Ligation between the 3’ end of the left stem arm and the 5’ end of the right stem arms result in “normal” gapped alignments. Ligation may occur at opposite ends of the stem, between the 5’ end of the left arm and the 3’ end of the right arm, resulting in chimeric alignments. The rearrangement of chimera performed in the earlier step ensured that all the alignments on the same strand and chromosome are represented by a single CIGAR string and one record (gap1.sam), while alignments with two arms on different strands or chromosomes are represented by two records each.

### 5.2 Network representation of non-continuous alignments

The reads/alignments are transformed into a network representation using the Python package NetworkX (Hagberg et al. 2008). Each vertex in the network represents a single alignment. Consider a pair of alignments *r_1_, r_2_*, each with left (*l*) and right (*r*) arm start and stop positions indexed by 0 and 1, respectively. Alignment *i* is represented by the 4-tuple of arm start and stop positions: (*a_i,l,0_, a_i,l,1_, a_i,r,0_, a_i,r,1_*). To determine whether the network vertices representing the alignment pair *r_1_, r_2_* are connected via an edge, left and right overlaps *o*_*j*_(*r_1_, r_2_*) and spans *s*_*j*_ (*r*_1_, *r*_2_), *j* ∈ {*l, r*} of the alignment pair are calculated. Overlap and span are defined as and are depicted in **Fig. 4**:

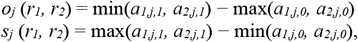

Note that the ratio of overlap to span is less than or equal to 0 if there is there is no overlap between arms, and is exactly 1 if the arms overlap completely and have the same length. An edge is drawn between the vertices representing alignments *r*_1_, *r*_2_ in the network if both left and right overlap ratios *o*_*j*_(*r_1_, r_2_*)/*s*_*j*_(*r_1_, r_2_*) exceed some overlap threshold *to*. This rule makes intuitive sense from a biological perspective: in order to have originated from the same stem structure, two alignments must have at least some overlapping portion in both arms. The sum of the right and left overlap ratios of alignments *r_1_, r_2_* is recorded as the weight of the edge connecting the vertices representing alignments.

To decrease graphing time, alignments are ordered by all four arm start and stop positions. For a given alignment ordering “primary” alignment— one half of a potential alignment pair—is selected, and a “secondary” alignment—the second half of a potential alignment pair—is drawn sequentially from the remaining ordered alignments. As long as the secondary alignment overlaps the primary alignment in both arms, the secondary alignment is added as a vertex to the network and the next alignment is selected as a candidate secondary alignment. However, once a candidate secondary alignment is found to share no overlap with the primary alignment, the remaining alignments in the ordering are skipped, and alignments from the next ordering are considered sequentially. This procedure of ordered traversals over all ordered alignments avoids an exhaustive pairwise search over non-overlapping alignments and increases pipeline efficiency. We refer to the resulting network comprising the set of vertices *V* and the set of edges *E* as the weighted alignments graph *G = (V, E)*.

### 5.3 Network clustering of non-continuous alignments

The graph comprising all alignments typically contains multiple subgraphs of connected components, which each represent groups of overlapping alignments that do not overlap with other groups (i.e., that overlap less than overlap threshold t_o_). However, the subgraphs themselves may sometimes be further broken down into distinct clusters of alignments that are more similar to each other than they are to other alignments in the subgraph. To account for this possibility, the CRSSANT pipeline first partitions the weighted alignments graph G into subgraphs of connected components, and then extracts clusters using two possible deterministic clustering methods: cliques-finding and spectral clustering.

Each method has its benefits and drawbacks. By definition, all vertices in a clique are fully connected, i.e. each vertex is accessible from every other vertex. In the biological setting, this can be interpreted as: all alignments whose network representations comprise a clique must overlap in both the left and right arms beyond threshold to. Intuitively, this implies that these alignments are all highly similar, and that the alignment arm start and stop positions have low variability. Similarly, the arm start and stop positions of the DG comprising these alignments can be obtained with high confidence. However, the requirement of full connectivity also has the undesirable potential to exclude a large number of alignments which could discard valuable sequencing information. Furthermore, the problem of finding a maximal clique is often very computationally demanding (Cazals and Karande 2008).

Spectral clustering, on the other hand, does not require full connectivity between all vertices assigned to the same cluster, and thus provides a more flexible method of grouping vertices in a graph. In addition, it is simple to implement, can typically be solved efficiently using modern software, and often outperforms simple clustering algorithms like k-means clustering (von Luxburg 2007). However, implementing spectral clustering depends on carefully choosing heuristics for a particular problem setting.

In the following subsections, we describe the two clustering methods and relevant parameters. The clustering parameter that is shared by both methods is the overlap threshold t_o_, which strongly affects subgraphs and, subsequently, the final DGs that alignments are clustered into using cliques-finding or spectral clustering. In general, a large to results in a larger number of subgraphs containing fewer alignments, since the alignments must overlap more before an edge is drawn between their respective nodes. Exactly how the number of subgraphs and their composition affects the outcome of each clustering method is discussed in each of the following subsections.

To speed up the clustering, we also implemented a sub sampling approach. Building a network representation and the subsequent clustering is computationally intensive when the number alignments is large. For each RNA, the number of new alternative conformations diminishes with more alignments. Once the alignment density reach a certain limit, we will not find any new conformations. Therefore, we limit the alignment density to a certain number, e.g. 1000, and all the additional alignments are added to the already established DGs after clustering. This approach is much faster than clustering all of them.

Clustering method 1: Cliques. This method, an adaptation of (Zhang et al. 2005), is built into the core NetworkX library and was used off-the-shelf. It divides each subgraph into cliques, or sets of nodes such that all nodes in the set share an edge with all other nodes. However, this implementation identifies all possible cliques that exist in the subgraph, meaning that a single alignment may exist in multiple cliques. This conflicts with our goal of matching each alignment to a single DG. To deal with this ambiguity, we filter the list of all possible cliques using a greedy approach: cliques are sorted in descending order based on the number of alignments in each clique, and candidate cliques are kept only if the set of kept cliques do not contain any of the alignments in the candidate clique. The final set of cliques that are kept are declared to be the DGs. This method tends to discard a large number of alignments, but has the benefit of resulting in DGs with highly similar alignments, i.e. that share significant overlaps in both arms.

Clustering method 2: Spectral clustering The spectral clustering method of cluster extraction is adapted from the Shi and Malik method described in (von Luxburg 2007), which we outline here briefly. For each subgraph *G*_*s*_ = *(V*_*s*_, *E*_*s*_) containing *m* > 1 vertices, we calculate the *m×m* dimensional degree matrix *D* whose components *d*_*i,j*_ are

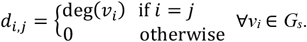

The degree deg(*v*_*i*_) of a vertex is the number of times an edge terminates at that vertex. In terms of the alignments from PARIS and related methods, deg(*v*_*i*_) represents the number of alignments overlapping the alignment represented by *v*_*i*_ with overlap ratio exceeding *t*_*o*_. We also calculate the *m×m* dimensional unnormalized Laplacian matrix *L* of *G*_*s*_ as *L = D − A*, where *A* is the adjacency matrix of *G*_*s*_ with components *A*_*i,j*_= **1**_*vi,vj*_, *a*_*i,j*_ ∈ {0, 1} indicating whether or not there exists an edge between vertices *v*_*i*_ and *v*_*j*_. In terms of the PARIS alignments, *A*_*i,j*_ indicates whether or not the sum of the overlap ratios of both arms of *vi* and *vj* exceed to. With *D* and *L*, we solve the generalized eigenproblem *Lv = λDv* using an eigengap heuristic based on the sorted eigenvalues and their corresponding eigenvectors to identify clustering parameter *k*, the number of clusters contained within the subgraph.

The eigengap heuristic chooses k such that the first k eigenvalues are “relatively small”, and the gap between the *k*^th^ eigenvalue *λk* and the (*k*+1)^th^ eigenvalue *λ*_k+1_ is “relatively large.” In practice, the variability of alignments assigned to the same subgraph can result in eigenvalues that are not well separated, making it hard to choose the first relatively large eigengap. This choice is further complicated by the presence of large eigengaps among larger eigenvalues, which disqualifies decision rules based on a global analysis of all eigenvalues. To address these challenges, our implementation of the eigengap heuristic considers only the first neig eigenvalues. Then, the *k*−1 eigengaps are calculated. For each eigengap we calculate what we call an “eigenratio,” i.e. the *i*^th^ eigenratio is the ratio between the *i*^th^ eigengap and the median of the preceding *i*−1 eigengaps. The first eigenratio is simply the first eigengap. Finally, *k* is determined to be one less than the index of the first eigenratio that exceeds an eigenratio threshold *t*_*eig*_.

Intuitively, the goal of clustering is to create groups such that the edges connecting different groups have very low weights, while the edges connecting vertices within a group have high weight. Note that the Laplacian matrix *L* may be thought of as quantifying all edges emanating “out” from vertices, which are exactly the components that are used as the basis of clustering. Thus, solving the generalized eigenproblem involving the Laplacian *L* and identifying the first *k* largest eigengaps may be interpreted as the solution to the principal components problem of identifying the k dimensions which provide optimal separation of vertex groups.

In practice, we observed that PARIS data tended to result in first eigengaps (the difference between the first and second eigenvalues) that were small, and whose ratios rarely exceeded *t*_*eig*_. This resulted in a preponderance of *k* = 0 or *k* ≥ 3, even in situations where a human arbiter would have decided on *k* = 2. To allow for the possibility of splitting subgraphs into just two DGs, we added a second component to the eigengap heuristic that first checks if the magnitude of the second eigenvalue is large (empirically, greater than 1), and then if the remaining *n*_*eig*_ − 2 eigenvalues are much smaller than the second eigenvalue (empirically, if the median of the remaining eigenvalues is at least an order of magnitude smaller than the magnitude of the second eigenvalue). If these conditions are met, then *k* is set to 2. Once *k* is chosen, we use the first k eigenvectors to perform k-means clustering (James et al. 2013) on the subgraph vertices. Each resulting cluster is a DG.

Spectral clustering parameters *n*_*eig*_ and *t*_*eig*_ have opposite effects on DG clustering. A larger number of candidate eigenvalues, *n*_*eig*_, results in more eigengaps considered during execution of the eigengap heuristic. Considering more eigengaps increases the possibility of choosing a larger *k*, which results in splitting the subgraph into a larger number of DG clusters, each containing fewer alignments. On the other hand, increasing the eigengap threshold teig tends to result in smaller *k* which, in turn, results in splitting the subgraph into a smaller number of DG clusters. Each of these clusters will contain more alignments than those that would have otherwise resulted from splitting the subgraph into a larger number of alignments based on a smaller *t*_*eig*_.

### 5.4 DG analysis

After alignments are grouped into DGs, the DGs are filtered to remove low-quality DGs, and to arrive at a final set of DGs. Singleton DGs containing only a single alignment are eliminated and the alignment is excluded from further analysis. This filtering criterion creates a direct relationship between the graph and clustering parameters t_o_, *n*_*eig*_ and *t*_*eig*_ and the final number of DGs and final number of alignment s included for subsequent analysis. When the parameters are chosen such that there are fewer alignments allocated to each DG, this increases the possibility that there exist singleton DGs, and in turn increases the possibility that more alignments are eliminated from later analysis steps.

In addition to eliminating singleton DGs and alignments, DGs are also checked for duplicate alignments. If any DG contains only alignments that are all identical in sequence, the alignments are deemed to be duplicates. The DG is again eliminated, and the duplicate alignments are excluded from further analysis. After DGs are filtered, various biologically significant attributes for each DG are calculated. The attributes are: number of alignments, arm start and stop positions, coverage and non-overlapping group.

Since the crosslink-ligation method produces one alignment per RNA stem structure, the number of alignments in each DG approximately corresponds to the cellular abundance of the stem structure represented by the DG. Potential DGs with very few alignments may either represent structures with low abundance, or they may be clusters of alignments that were incorrectly mapped due to sequencing errors. Thus, the number of alignments making up a DG are recorded as an important attribute.

The 4-tuple of DG arm start and stop positions are calculated as the median of arm start and stop positions of all alignments in the DG, and are an approximation of the gene sequence underlying the alignments that engages in stem structure formation. The relative coverage of DG *i* is defined in (Lu et al. 2016) to be 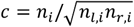 where ni is the total number of alignments in DG *i*, *nl,i* is the number of alignments whose left arms overlap the left arm of DG *i* and *nr,i* is the number of alignments whose right arms overlap the right arm of DG *i*. Coverage is a score with range (0, 1], and is an approximation of how much the genomic region spanned by a DG engages in the formation of multiple RNA structures. A coverage score equals to the maximum value of 1 indicates that the genomic region spanned by the DG gives rise to only one RNA stem structure. On the other hand, a low coverage close to zero suggests that the genomic region spanned by the DG produces numerous RNA stem structures, in other words, that the genomic region engages in formation of multiple alternative RNA stem structures.

### 5.5 NG clustering

Finally, for each DG a non-overlapping group (NG) is calculated. DGs are assigned to NGs such that there are no overlaps between any of the alignments in all DGs assigned to a given NG. NG assignments allow DGs to be easily viewed in genomic visualization software, like the popular Interactive Genomics Viewer (Robinson et al. 2011).

### 5.6 Output files from CRSSANT.py

The CRSSANT pipeline analyzes alignments by gene pairs and produces two types of output files for alignments overlapping a given gene pair (*g1, g2*). For each gene pair, only alignments from the input SAM file whose left arms map to *g1* and whose right arms map to *g2* are analyzed. If alignments from the gene pair form valid DGs that do not contain any non-overlapping alignments, the two output files are produced: a SAM file and a BEDPE file. It is important to check for nonoverlapping alignments within the same DG, since the presence of an alignment left arm begins downstream of other alignments’ right arms in the same DG is invalid—there is no biological basis for such alignments being clustered together and no evidence that they originated in the same genomic region. As a result, if any DGs contain non-overlapping alignments, DG results are not written to output files.

All alignments successfully clustered into DGs that passed the filtering step are written to the output SAM file. This SAM file is identical to the input SAM file, with two additional annotations for DGs and NGs. For each alignment assigned to DG *N*, the string DG:i:g1_g2_N is appended to the alignment line in the SAM file. The string NG:i:M is also appended to the alignment line, where *M* is the NG to which DG N belongs. Attributes of these DGs are written to the output BEDPE file which includes the genomic region of the DG, the DG identification number, coverage, number of alignments, arm start and stop positions and arm lengths.

## 6. Analysis of U2 snRNA structures

Human U2 snRNA sequence and conformations ATCGCTTCTCGGCCTTTTGGCTAAGATCAAGTGTAGTATCTGTTCTTATCAGTTTAATATCTGATACGTCCTCTATCCGAGGACAATATATTAAATGGATTTTTGGAAATAGGAGATGGAATAGGAGCTTGCTCCGTCCACTCCACGCATCGACCTGGTATTGCAGTACTTCCAGGAACGGTGCACC Conformation SLIIa and SLIIb, plus the SLIId blocking the branch point recognition sequence (BPRS):......(((.((((....)))).))).......((((((.......((((((........)))))).((((((.....))))))...))))))..................((((.((((...((((....)))).))))))))..((((((.(((((.............)))))..))))))... Conformation SLIIb and SLIIc......(((.((((....)))).))).........................(((((((((.......((((((.....))))))...)))))))))...............((((.((((...((((....)))).))))))))..((((((.(((((.............)))))..))))))...

Alternative conformations at SLIII and SLIV.

GGAAAUAGGAGAUGGAAUAGGAGCUUGCUCCGUCCACUCCACGCAUCGACCUGGUAUUGCAGUACUUCCAGGAACGGUGCACC.......((((.((((...((((....)))).))))))))..((((((.(((((.............)))))..))))))...............(((((..(..((..................................))..)..))))).............

The following are manually curated references for mapping human and mouse snRNAs.

>hssnRNA

ATACTTACCTGGCAGGGGAGATACCATGATCACGAAGGTGGTTTTCCCAGGGCGAGGCTTATCCATTGCACTCCGGATGTGCTGACCCCTGCGATTTCCCCAAATGTGGGAAACTCGACTGCATAATTTGTGGTAGTGGGGGACTGCGTTCGCGCTTTCCCCTGNNNNNNNNNNNNNNNNNNNNNNNNNNNNNNNNNNNNNNNNNNNNNNNNNNNNNNNNNNNNNNNNNNNNNNNNNNNNNNNNNNNNNNNNNNNNNNNNNNNNATCGCTTCTCGGCCTTTTGGCTAAGATCAAGTGTAGTATCTGTTCTTATCAGTTTAATATCTGATACGTCCTCTATCCGAGGACAATATATTAAATGGATTTTTGGAAATAGGAGATGGAATAGGAGCTTGCTCCGTCCACTCCACGCATCGACCTGGTATTGCAGTACTTCCAGGAACGGTGCACCNNNNNNNNNNNNNNNNNNNNNNNNNNNNNNNNNNNNNNNNNNNNNNNNNNNNNNNNNNNNNNNNNNNNNNNNNNNNNNNNNNNNNNNNNNNNNNNNNNNNAGCTTTGCGCAGTGGCAGTATCGTAGCCAATGAGGTTTATCCGAGGCGCGATTATTGCTAATTGAAAACTTTTCCCAATACCCCGCCGTGACGACTTGCAATATAGTCGGCATTGGCAATTTTTGACAGTCTCTACGGAGACTGGNNNNNNNNNNNNNNNNNNNNNNNNNNNNNNNNNNNNNNNNNNNNNNNNNNNNNNNNNNNNNNNNNNNNNNNNNNNNNNNNNNNNNNNNNNNNNNNNNNNNGTGCTCGCTTCGGCAGCACATATACTAAAATTGGAACGATACAGAGAAGATTAGCATGGCCCCTGCGCAAGGATGACACGCAAATTCGTGAAGCGTTCCATATTTTNNNNNNNNNNNNNNNNNNNNNNNNNNNNNNNNNNNNNNNNNNNNNNNNNNNNNNNNNNNNNNNNNNNNNNNNNNNNNNNNNNNNNNNNNNNNNNNNNNNNATACTCTGGTTTCTCTTCAGATCGCATAAATCTTTCGCCTTTCATCAAAGATTTCCGTGGAGAGGAACAACTCTGAGTCTTAACCCAATTTTTTGAGCCTTGCCTTGGCAAGGCTANNNNNNNNNNNNNNNNNNNNNNNNNNNNNNNNNNNNNNNNNNNNNNNNNNNNNNNNNNNNNNNNNNNNNNNNNNNNNNNNNNNNNNNNNNNNNNNNNNNNAAAAAGGGCTTCTGTCGTGAGTGGCACACGTAGGGCAACTCGATTGCTCTGCGTGCGGAATCGACATCAAGAGATTTCGGAAGCATAATTTTTTGGTATTTGGGCAGCTGGTGATCGTTGGTCCCGGCGCCCTTTNNNNNNNNNNNNNNNNNNNNNNNNNNNNNNNNNNNNNNNNNNNNNNNNNNNNNNNNNNNNNNNNNNNNNNNNNNNNNNNNNNNNNNNNNNNNNNNNNNNNATGCCTTAAACTTATGAGTAAGGAAAATAACGATTCGGGGTGACGCCCGAATCCTCACTGCTAATGTGAGACGAATTTTTGAGCGGGTAAAGGTCGCCCTCAAGGTGACCCGCCTACTTTGCGGGATGCCTGGGAGTTGCGATCTGCCCGNNNNNNNNNNNNNNNNNNNNNNNNNNNNNNNNNNNNNNNNNNNNNNNNNNNNNNNNNNNNNNNNNNNNNNNNNNNNNNNNNNNNNNNNNNNNNNNNNNNNAACCATCCTTTTCTTGGGGTTGCGCTACTGTCCAATGAGCGCATAGTGAGGGCAGTACTGCTAACGCCTGAACAACACACCCGCATCAACTAGAGCTTTTGCTTTATTTTGGTGCAATTTTTGGAAAAATNNNNNNNNNNNNNNNNNNNNNNNNNNNNNNNNNNNNNNNNNNNNNNNNNNNNNNNNNNNNNNNNNNNNNNNNNNNNNNNNNNNNNNNNNNNNNNNNNNNNGTGTTGTATGAAAGGAGAGAAGGTTAGCACTCCCCTTGACAAGGATGGAAGAGGCCCTCGGGCCTGACAACACGCATACGGTTAAGGCATTGCCACCTACTTCGTGGCATCTAACCATCGTTTTT

>mmsnRNA

ATACTTACCTGGCAGGGGAGATACCATGATCACGAAGGTGGTTTTCCCAGGGCGAGGCTTATCCATTGCACTCCGGATGTGCTGACCCCTGCGATTTCCCCAAATGCGGGAAACTCGACTGCATAATTTGTGGTAGTGGGGGACTGCGTTCGCGCTCTCCCCTGNNNNNNNNNNNNNNNNNNNNNNNNNNNNNNNNNNNNNNNNNNNNNNNNNNNNNNNNNNNNNNNNNNNNNNNNNNNNNNNNNNNNNNNNNNNNNNNNNNNNATCGCTTCTCGGCCTTTTGGCTAAGATCAAGTGTAGTATCTGTTCTTATCAGTTTAATATCTGATACGTCCTCTATCCGAGGACAATATATTAAATGGATTTTTGGAACTAGGAGTTGGAATAGGAGCTTGCTCCGTCCACTCCACGCATCGACCTGGTATTGCAGTACCTCCAGGAACGGTGCAACNNNNNNNNNNNNNNNNNNNNNNNNNNNNNNNNNNNNNNNNNNNNNNNNNNNNNNNNNNNNNNNNNNNNNNNNNNNNNNNNNNNNNNNNNNNNNNNNNNNNAGCTTTGCGCAGTGGCAGTATCGTAGCCAATGAGGTTTATCCGAGGCGCGATTATTGCTAATTGAAAACTTTTCCCAATACCCCGCCGTGACGACTTGCAATATAGTCGGCATTGGCAATTTTTGACAGTCTCTACGGAGACTGGNNNNNNNNNNNNNNNNNNNNNNNNNNNNNNNNNNNNNNNNNNNNNNNNNNNNNNNNNNNNNNNNNNNNNNNNNNNNNNNNNNNNNNNNNNNNNNNNNNNNGTGCTCGCTTCGGCAGCACATATACTAAAATTGGAACGATACAGAGAAGATTAGCATGGCCCCTGCGCAAGGATGACACGCAAATTCGTGAAGCGTTCCATATTTTNNNNNNNNNNNNNNNNNNNNNNNNNNNNNNNNNNNNNNNNNNNNNNNNNNNNNNNNNNNNNNNNNNNNNNNNNNNNNNNNNNNNNNNNNNNNNNNNNNNNATACTCTGGTTTCTCTTCAGATCGTATAAATCTTTCGCCTTTTACTAAAGATTTCCGTGGAGAGGAACAAATCTGAGTCTTAACCCAATTTTTTGAGGTCTTGTGCTTACAAGACTNNNNNNNNNNNNNNNNNNNNNNNNNNNNNNNNNNNNNNNNNNNNNNNNNNNNNNNNNNNNNNNNNNNNNNNNNNNNNNNNNNNNNNNNNNNNNNNNNNNNAAAAAGGGCTTCTGTCGTGAGTGGCACACGCAGGGCAACTCGATTGCTGTGCGTGCGGAATCGACATCAAGAGATTTCGGAAGCATAATTTTTTGGTAATTGGGCAGCTGGTGATCGTTGGTCCCGGCGCCCTTGNNNNNNNNNNNNNNNNNNNNNNNNNNNNNNNNNNNNNNNNNNNNNNNNNNNNNNNNNNNNNNNNNNNNNNNNNNNNNNNNNNNNNNNNNNNNNNNNNNNNATGCCTTAAACTTATGAGTAAGGAAAATAACGATTCGGGGTGACGCCCGAGTCCTCACTGCTTATGTGAGAAGAATTTTTGAGCGGGTATAGGTTGCAATCTGAGGCGACCCGCCTACTTTGCGGGATGCCTGGGTGACGCGATCTGCCCGNNNNNNNNNNNNNNNNNNNNNNNNNNNNNNNNNNNNNNNNNNNNNNNNNNNNNNNNNNNNNNNNNNNNNNNNNNNNNNNNNNNNNNNNNNNNNNNNNNNAACCATCCTTTTCTTGGGGTTGCGCTACTGTCCAATGAACGCGTAGTGAGGGCAGTACTGCTAACGCCTGACAACACACCTGCATCGGTTAGAGCTCTGCTTTACCTTGGTGCAATTTTTGGAAAAAANNNNNNNNNNNNNNNNNNNNNNNNNNNNNNNNNNNNNNNNNNNNNNNNNNNNNNNNNNNNNNNNNNNNNNNNNNNNNNNNNNNNNNNNNNNNNNNNNNNNNNGTGTTGTATGAAAGGAGAGAAGGTTAGCACTCCCCTTGACAAGGATGGAAGAGGCCCTTGGGCCTAACAACACACATACGGTTAAGGCATTGCCACCTACTTCGTGGCATCTAACCACTGTTTTT

## 7. Identification and structure analysis of U3 genes

The U3 snoRNA has multiple paralogs in the mammalian genome. To make the human U3 reference to represent the most abundant paralog, we examined the human genome U3 expression in HEK293 PARIS data mapped to hg38. The human genome encodes 5 U3 genes for which there is detectable expression, SNORD3A, SNORD3B-1, SNORD3B-2, SNORD3C and SNORD3D (**Supplemental Table 3**, coverage refers to max depth). The first 5 isoforms A-D are located in a small cluster on chromosome 17 (chr17:19061900-19190300), while the other paralogs are scattered in the genome. The two SNORD3B genes are identical in sequence. They only have 4 single nucleotide differences among all of them. Based on these numbers, we selected SNORD3C as the representative human U3 gene and named it hsU3 for making the STAR index and IGV reference (Robinson et al. 2011).

To make the mouse U3 reference that represents the most abundant paralog if multiple paralogs are expressed, we used Rfam U3 sequence as query and performed BLAST (discontiguous megablast) search for paralogous sequences in mouse genomic + transcript (Mouse G+T) with the following parameters that are different from default: unchecking “Low complexity regions and “Species-specific repeats”. The standard parameters are as follows: max target sequences: 100, short queries: Automatically adjust parameters for short input sequences, Expect threshold: 10, word size: 11, max matches in a query range: 0, match/mismatch scores: 2,-3, gap costs: existence 5, extension 2, mask for lookup table only: checked. The BLAST search returned 10 sequences in the mouse genome with >90% coverage of the query. To identify the expressed paralogs, we checked the PARIS data from mES cells (**Supplemental Table 3**). The three most abundant paralogs Rnu3b1, Rnu3b3 and Rnu3b4 are identifical and therefore used as the reference. The mouse U3 is named mmU3 for making the STAR index and IGV reference. The indexing and mapping parameters are the same as for the human PARIS data.

Structure models for human U3 were built based on thermodynamics and PARIS as follows.

>hsU3

AAGACTATACTTTCAGGGATCATTTCTATAGTGTGTTA CTAGAGAAGTTTCTCTGAACGTGTAGAGCACCGAAAAC CACGAGGAAGAGAGGTAGCGTTTTCTCCTGAGCGTGAA GCCGGCTTTCTGGCGTTGCTTGGCTGCAACTGCCGTCA GCCATTGATGATCGTTCTTCTCTCCGTATTGGGGAGTG AGAGGGAGAGAACGCGGTCTGAGTGGT....((((((.((((((((..((((((.(((((...))))).)))))).)))))))).))))))..........(((((..(((.........(((( ((((((((((.(((.((...((((..(.((((((((......))))).))))..)))).)).))).)))(((((((((((.....)))))).))))))))))))))))..)))..)))))

**Supplemental Table 3.** Human and mouse U3 genes.

| Human U3 | coverage | Mouse U3 (mm10) | coverage |
| --- | --- | --- | --- |
| SNORD3A | 56 | chr1:39519533-39519746 | 92 |
| SNORD3B-1 | 205 | chr10:40258467-40258679 | 552 |
| SNORD3B-2 | 12 | chr11:87443159-87443530 | 3589 (Rnu3b1) |
| SNORD3C | 513 | chr11:87471290-87471661 | 73 (Rnu3b2) |
| SNORD3D | 344 | chr11:87448549-87448920 | 4865 (Rnu3b3) |
| SNORD3E | 5 | chr11:87462209-87462580 | 2818 (Rnu3b4) |
| SNORD3F | 1 | chr11:96032678-96032891 | 55 |
| SNORD3G-J | 0 | chr19:4078281-4078528 | 170 |
| SNORD3K | 1 | chr12:102253685-102253474 | 328 |
| SNORD3P3 | 4 | chr2:121004089-121004301 | 50 |

## 8. Identification and structure analysis of U8 genes

Four close paralogs of human U8 can be identified in the genome by BLAST (top 7 loci shown in **Supplemental Table 4**) and inspection of gene expression based on published high throughput RNA-seq (MCF7 cells and K562 cells, GSE88622 and GSE78663) and human HEK293 PARIS data (Lu et al. 2016) showed only the one located in the 3’UTR of TMEM107 is expressed (chr17:8173453-8173588). This is consistent with the human genetics study showing that mutations in this U8 gene cause LCC (Jenkinson et al. 2016). The Rfam sequence for human U8 contains several nucleotide variants compared to the TMEM107 and it is incorrect. The first gene locus below is the SNORD118 (chr17: 8173453-8173588). The human U8 sequence and two alternative structure models are as follows: ATCGTCAGGTGGGATAATCCTTACCTGTTCCTCCTCCGGAGGGCAGATTAGAACAT GATGATTGGAGATGCATGAAACGTGATTAACGTCTCTGCGTAATCAGGACTTGCAACACCCTGATTGCTCCTGTCTGATT.....((((((((......)))))))).((((((...))))))..(((((((.((........(((((((((.((.((((.....)))).))))))) (((((((...........))))))).)))))))))))))...((((((((((((((((.(((..(((((((((...)))))).........)))))).)))))...(((((.((.((((.....)))).))))))) (((((((...........)))))))..))))))))))).

**Supplemental Table 4.**
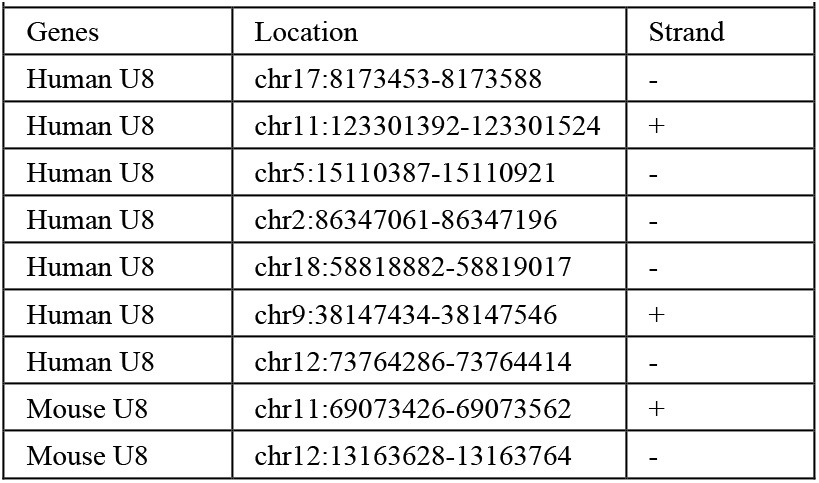
Human and mouse U8 genes.

| Genes | Location | Strand |
| --- | --- | --- |
| Human U8 | chr17:8173453-8173588 | - |
| Human U8 | chr11:123301392-123301524 | + |
| Human U8 | chr5:15110387-15110921 | - |
| Human U8 | chr2:86347061-86347196 | - |
| Human U8 | chr18:58818882-58819017 | - |
| Human U8 | chr9:38147434-38147546 | + |
| Human U8 | chr12:73764286-73764414 | - |
| Mouse U8 | chr11:69073426-69073562 | + |
| Mouse U8 | chr12:13163628-13163764 | - |

Using the Rfam mouse U8 as a query, we searched the mouse genome for U8 paralogs using BLAST and results are as follows (**Supplementary Table 4**). The closest sequence was mapped to the chr11:69073426-69073562, near the 3’ end of mouse Tmem107 (just like the human SNORD118 next to TMEM107). The genomic copy has a 1nt insertion likely due to genome sequencing errors.

Inspection of mouse E14 and G1E cell line RNA-seq data (GSE90277 and GSE101173) and mouse PARIS data confirmed that this U8 gene is actively expressed and the 1nt insertion (Lu et al. 2016). The start and end of this transcript was also verified. The sequence is 136nt long, shown as follows, together with the two structure conformations in dot-bracket format based on thermodynamics and PARIS.

ATCGTCAGGAGGTTAATCCTTACCTGTTCCTCCTTCGGAGGGCAGTAGAAAATGATGATTGGAGCTTGCATGATCTGCTGATTAGCATTTCCATGCAATCAGGACCTGACAACATCCTGGTTGCTCCTATCTGATT...(((((((((...........((((.(((((...)))))))))..............(((((..(((.(((((....)))))))).))))).((( (((((((..((....)))))))))))).))).)))))).

...((.((((......)))).))((((.(((((...))))))))).....(((.(.((((((((..(((.(((((....)))))))).))))).((( (((((((..((....))))))))))))....)))).)))

## 9. Identification and structure analysis of U13 genes

Tyc and Steitz first described U3, U8 and U13 in 1989 (Tyc and Steitz 1989). The estimated abundance for these snoRNAs are as follows: 200000, 40000 and 10000 copies per cell. In this original report, the U13 snoRNA was annotated to be 104nt, but it was not accurate. Here, human U13 (SNORD13) genes were identified by BLAST against the human genome + transcriptome using the sequence from Tyc and Steitz (Tyc and Steitz 1989) using these parameters that are different from default: unchecking “Low complexity regions and “Species-specific repeats” (**Supplemental Table 5**, top hits out of 25 are listed). The standard parameters are as follows: max target sequences: 100, short queries: Automatically adjust parameters for short input sequences, Expect threshold: 10, word size: 11, max matches in a query range: 0, match/mismatch scores: 2,-3, gap costs: existence 5, extension 2, mask for lookup table only: checked.

Visual inspection of human HEK293 PARIS data revealed a single location with significant mapped reads, at chr8:33513475-33513578 in hg38, upstream of the TTI2 gene, suggesting that this is the functional gene copy. All the other loci have no mapped reads. The coverage of reads extends over the 3’ end of the annotated location, suggesting the original annotation was not the full length. Similarly, the mouse U13 homolog was identified, and the coverage of reads extends over the 3’ end of the annotated location, confirming that the previous annotation was not full length. By comparing human and mouse sequences beyond the 3’ end, 15 nts were added to the human U13, while the mouse U13 was extended 16nt.

**Supplemental Table 5.** Human and mouse U13 genes.

| Genes | Location | Highest coverage |
| --- | --- | --- |
| Human U13 | chr8:33513475-33513577 | 330 (few mismatches) |
| Human U13 | chr8:101847257-101847342 | 0 |
| Human U13 | chr8:58662247-58662344 | 0 |
| Human U13 | chr8:94906017-94906119 | 2 |
| Human U13 | chr2:172691402-172691299 | 0 |
| Human U13 | chr3:47250525-47250626 | 39 (only in middle) |
| Human U13 | chr3:179614775-179614878 | 1 |
| Human U13 | chr16:53334562-53334662 | 14 |
| Human U13 | chr18:2774882-2774962 | 1 |
| Mouse U13 | chr8:31149864-31149967 | 900 (no mismatches) |
| Mouse U13 | chr6:38057974-38058076 | 89 (only in middle) |
| Mouse U13 | chr4:132485885-132485994 | 0 |
| Mouse U13 | chr1:57865751-57865840 | 0 |

BLAST search in the mouse genome revealed four paralogs: one in chr8:31149803-31150056, the other three on chr1, chr4 and chr6. The numbers of reads mapped to these locations clearly show that the one on chr8 is a real gene, while the others are pseudogenes The revised human U13 sequence is 120nt long and the revised mouse U13 is 121nt long. Their predicted secondary structures are as follows. Extension at the 3’ end allows the formation of terminal stemloops that connects the 5’ and the 3’ end.

>hsU13

GATCCTTTTGTAGTTCATGAGCGTGATGATTGGGTGTTCATACGCTTGTGTGAGATGTGCCACCCTTGAACCTTGTTACGACGTGGGCACATTACCCGTCTGACCTGAACTTCAAGGATC ((((((..((.(((((..(((.((.......((((((((((((....)))))))......)))))....)))))((((.((((.((.))))))))))..))))).))))))))

>mmU13

GATCCTTTCTGGTTCATAAAGCGTGATGATTGGGTGTTCACGCCATTGCGTGACATGTGCCGCCCATAAACCTTGTTACGACGTGGGCACATTACCCGTCTGACATGAACTTAGGAGGATC ((((((((..(((((((.(((.((......((((((.((((((....)))))).......))))))...)))))((((.((((.((.)))))))))))))))))..))))))))

To identify more distant U13 paralogs, significant revision of the model and analysis of alignments is needed. We examined the alignments from Rfam and focused on several sequences outside of the mammals. Ixodes U13 was already identified from Rfam. Yeast snR45 is likely the U13 homolog based on previous analysis (Sharma et al. 2017). Plants also seem to have U13 homologs (Kim et al. 2010). The alignments of human, yeast and plant U13 snoRNAs were presented in Sharma et al. 2017 (Plos Genetics). However, the alignments seem to be missing the 3’ end, just like the human and mouse U13 annotations. The extended gene models and secondary structures are presented below.

>Ixodes U13

<u>ATTCTTCCGAGTTTCA</u>ATGGGGAGTGATGAGTCGTGGGTGTTCATGCCAATGCGTGATACGTGCGAC<u>GCCCTGATTGTTACGAC</u>GATTGCACGTGAC CCCTCTGACCCCATTGGCGGAAC....(((((....(((((((((........((((((.((((.(((((....))))))))).)))))).........((((((.........))))))........)))))))))))))).

3’ end of Ixodes 18S rRNA and internal structure. The two regions that base pair with U13 are labeled with lines above the sequence:

GAGGAAGTAAAAGTCGTAACAAGGTTTCCGTAGGTGAACCTGCGGAAGGATCATTA........................((((((((((....))))))))))........

S. cerevisiae snR45: <u>ATGACCTTCCAAGTTTT</u>TAAAAGAATACGATGATATTATTTGCGTTTCAAATCGAACAATTCTTCTCGGAGCGATCTGAGGTTTTAATGGAGATAGC GGTTCCTGCGCAACCCATT<u>GATCTTGTTAC</u>ATTCTTAAGAATGACAAGGACGCTTTTATAAAATTCTGATTCTTT...................(((((((..((...........(((((.................(((((((....)))))))..(((((((.....((((...)))).....))))))).(((((..(((((....)))))))))))))))...........))..)))))))

Here the left side is the 3’ end of Ixodes 18S, and the right side is the Ixodes U13, where N corresponds an omitted region. The total number of base pairs is 13 + 15 = 28, close to the structural model for human 18S vs. U13 (15 + 17 = 32). The start and end of the base pairing regions for both Ixodes 18S and U13 are the same as for human, further supporting that this is the homologous interaction.

AAAGUCGUAACAAGGUUUCCGUAGGUGAACCUGCGGAAGGAUCAUUA&AUUCUUCCGAGUUUCAAUNACGCCCUGAUUGUUACGACGA...(((((((((((((.........((((.((.(((((((((......))))))))))).)))).....)))....))))))))))..

As a comparison, here is the human 18S:U13 interaction: AAAGUCGUAACAAGGUUUCCGUAGGUGAACCUGCGGAAGGAUCAUUA&GAUCCUUUUGUAGUUCAU&CACCCUUGAACCUUGUUACGAC...(((((((((((((((......((((((.((((((((((((....&)))))))))))))))))).)))))..)))))))))))))))

---

**Supplemental Figure 1.**
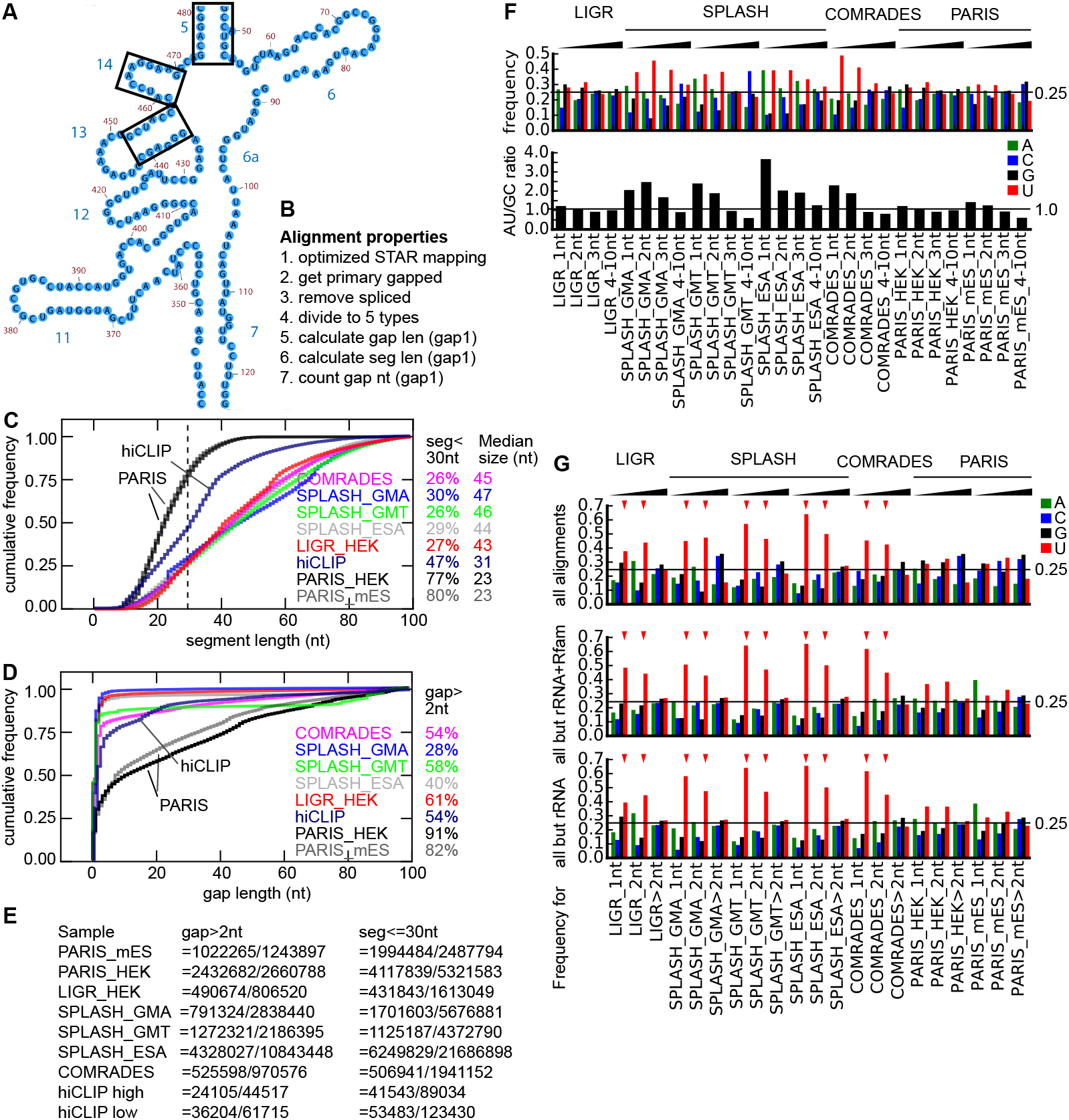
Analysis of segment length, gap length and gap nucleotides. (A) Part of the 5’ domain in human 18S rRNA (47-479nt), helices 5-14 (adapted from the gatech ribosome gallery). The two arms of h5 are 6 and 7 nucleotides respectively, while the two arms of h14 are only 4nt each. (B) Outline of the procedure for the analysis of segment length, gap length and gap nucleotide properties. Reads were first mapped to the genome reference using optimized permissive STAR parameters, and primary alignments were extracted for analysis. Spliced alignments were removed. Then the remaining alignments were divided into 5 categories (excluding the bad.sam). Segment and gap properties were calculated for the gap1.sam output. These datasets were used: PARIS (HEK1 bc07), PARIS (mES), LIGR (SRR3361013 HEK), SPLASH_GMA (SRR3404937, GM12892, polyA), SPLASH_GMT (SRR3404942, GM12892 total), SPLASH_ESA (hES), COMRADES (ZIKV) and all hiCLIP. (C) Cumulative distributions of segment lengths for gap1.sam for data from various published methods. (D) Cumulative distributions of gap lengths for gap1.sam alignments from various published methods. (E) Alignment numbers for each dataset with gaps >2nt and segment lengths <=30nt (over total numbers of reads or segments). (F-G) Small gaps are primarily U deletions, likely due to psoralen adducts. Nucleotide compositions were calculated for 1, 2, 3 and 4-10nt gaps separately for alignments mapped to hg38 or mm10 (F), or 1, 2 and >2nts for alignments mapped to Rfam+mRNA reference (G). In panel (G), alignments were further categorized in 3 subsets to show the gap nucleotide properties: all alignments (top), all alignments except rRNA+Rfam (middle), and all alignments except rRNA (bottom).

**Supplemental Figure 2.**
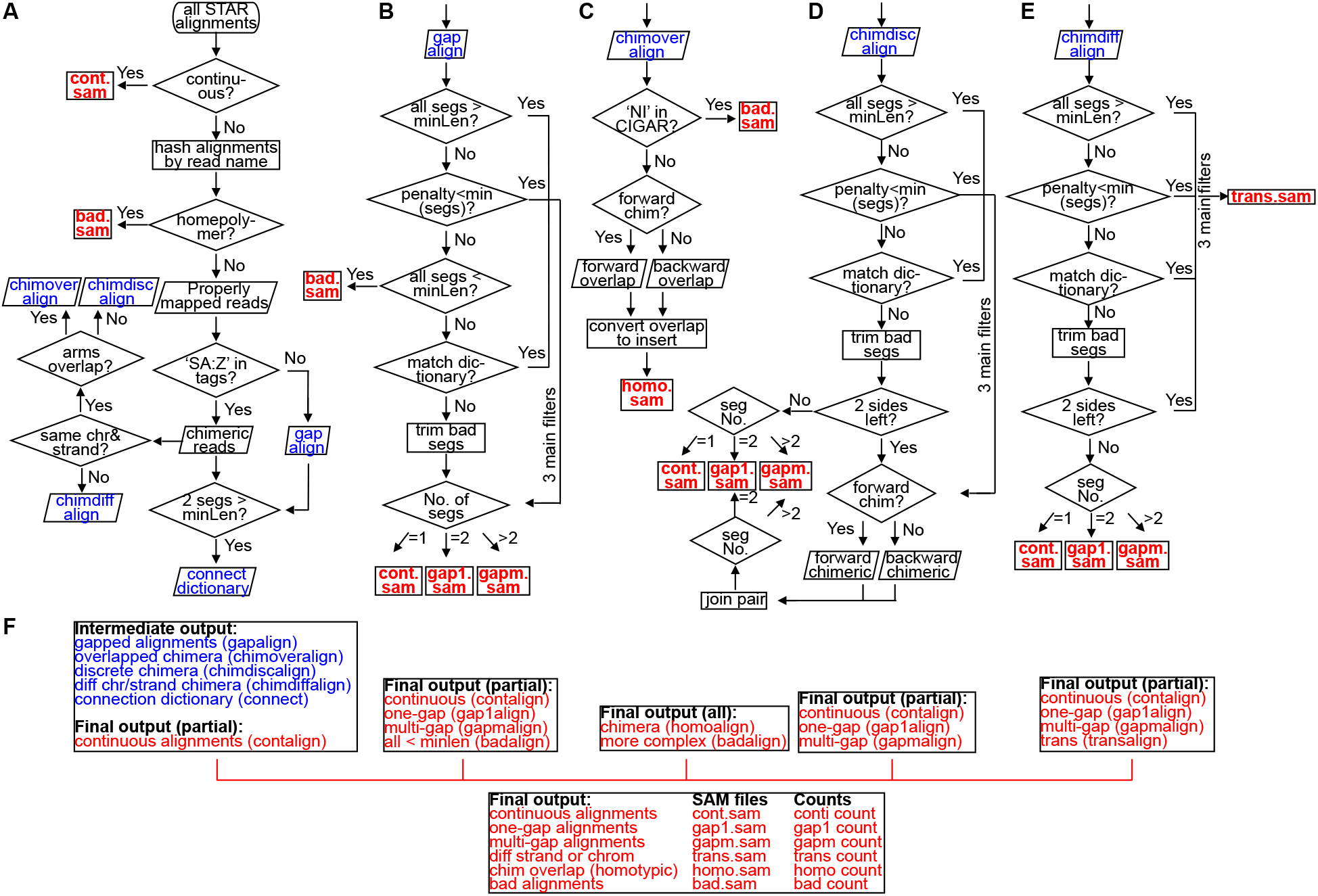
Design of the algorithms for processing non-continuous alignments from crosslink-ligation methods (gaptypes.py). Blue: intermediate data and files. Red: final output data and files. (A) All reads are first divided to different categories and the non-continuous alignments with segments >minlen (minimal length, e.g. 15) are used to build a connection dictionary, similar to the splice junction database. Note, reads and alignments are different by definition. Here all alignments were hashed using read names. One read may correspond to one or multiple alignments. It is important to point out that “reference” or “chromosome” are artificially defined; they are dif-ferent from the typical uses in biology. (B-E) Processing flowcharts for each type of alignment. This design has three criteria in filtering. (1). If all fragments are >minLen (e.g. 15nt), keep, otherwise go to step 2. (2). If all the fragments are “close” to the long matches (>=15), keep, otherwise go to step 3. (3). If all fragments overlap existing “good” alignments, keep, otherwise discard the short segments. (F) Summary of the output data and formats from each processing stage and the final output.

**Supplemental Figure 3.**
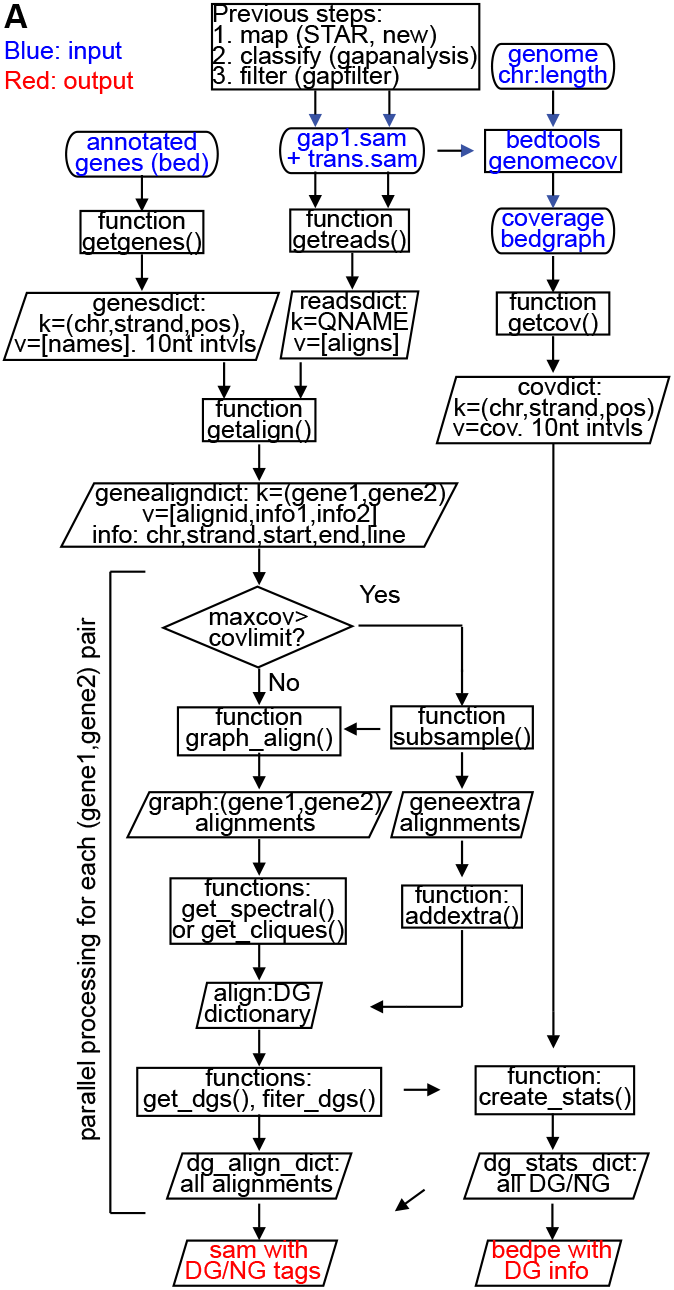
Flowchart for the CRSSANT method. (A) Blue: input data and files. Red: output data and files. Input data are annotated genes (bed format), gap1 and trans alignments (sam format), and genome length information file (chr \t length format). Annotated genes are first converted to a dictionary (hash table in python) with key (chr, strand, pos) and value of gene names overlapping the key position in the genome. Positions are in 10nt intervals, e.g. (chr1, ‘+’, 0), (chr1, ‘+’, 10), etc. Alignments from gap1 and trans files were first converted to a reads dictionary, where key is read name (QNAME) and value is a list of its alignments to the genome. The coverage bedgraph was calculated from input sam files and genome information file. Then the coverage bedgraph was converted to a coverage dictionary (covdict), where key is (chr, strand, pos) and value is coverage at the key position. The getalign() function merges the genesdict and readsdict to a dictionary to match alignments to gene pairs, where the key is a pair of genes (gene1, gene2), and value is (alignid, infor1, info2). Here gene1 and gene2 can be either the same (intramolecular structures) or different (intermolecular interactions). The following steps are parallelized to speed up the processing (depending on computer cpu numbers). The subsample() function sub-samples alignments to speed up processing for regions where coverage is too high (e.g. sub-samples alignments when coverage exceeds covlimit). The extra alignments not sub-sampled are added back to the DGs once an initial set of DGs are generated. Therefore, all alignments are used in the final output. Output files including a sam file and a bedpe file. In the sam file, each alignment has a DG (duplex group) tag and an NG (non-overlapping group) tag. In the bedpe file, detailed information of the DGs is collected.

**Supplemental Figure 4.**
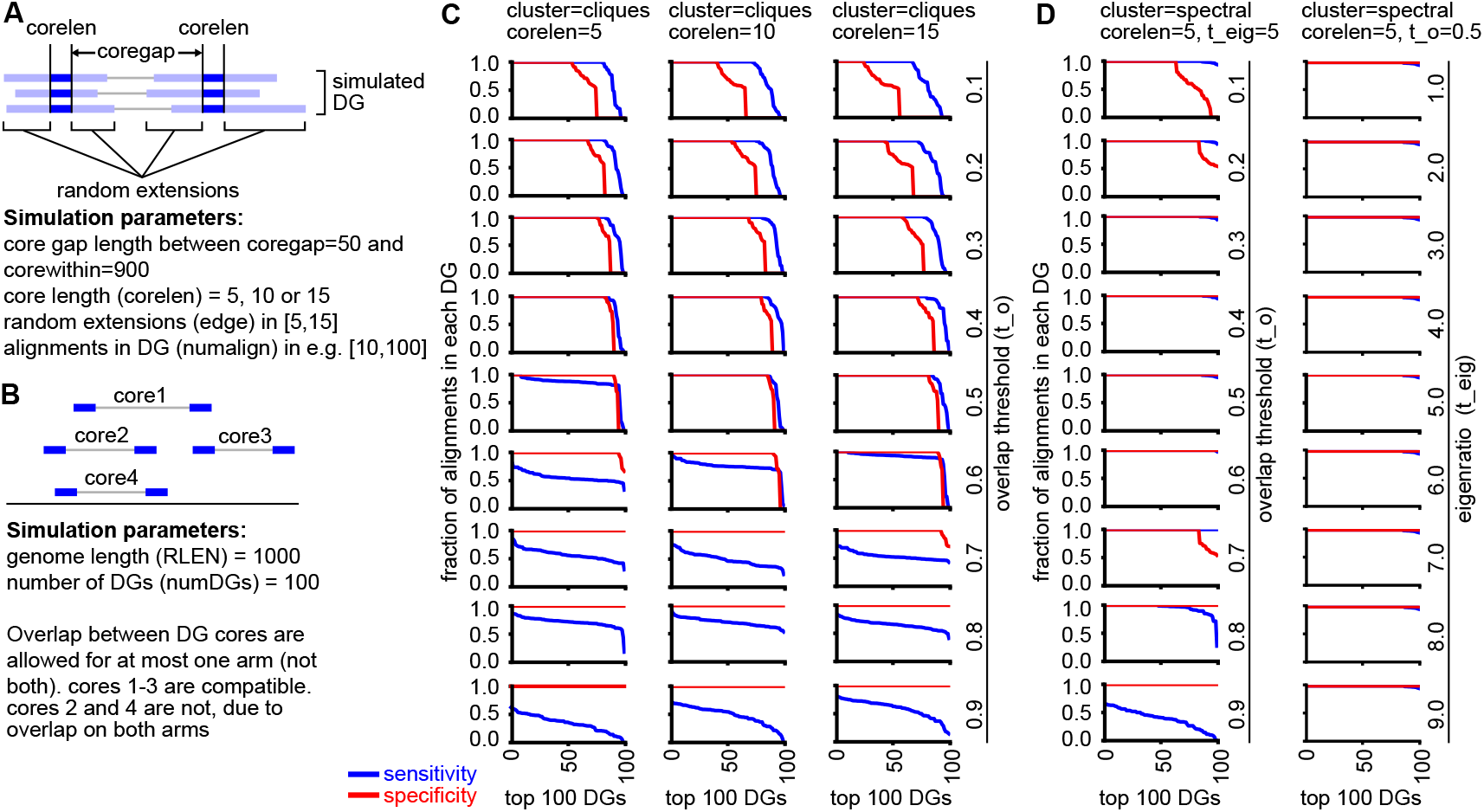
Benchmarking CRSSANT with simulated DGs. (A-B) Design of the DG simulation algorithm (dgsim.py). First, on an artificial chromosome of chr1:1-1000, made of nucleotides “N” in one gene GENE1: 1-1000, pairs of core intervals of a specified length (corelen, e.g. 5, 10 or 15nts) are selected, and randomly positioned in the range of chr1:100-900. The first 10kb of hg38 chr1 happens to be a stretch of “N”, so the results can be viewed on hg38. The two intervals of each pair are at least coregap away from each other (e.g. coregap=50), and within a specific distance (e.g. corewithin=1000, for chr1:1-1000). Each side of the two cores are extended by a random length (e.g. in the range [5,15]) to make one gap1 alignment. Each pair of core intervals are expanded to a set number of alignments that make up one DG, and the number of alignments in each DG is randomly set in a specific range, e.g. DGlower=10, DGupper=100. A set number of DGs are generated (e.g. 100), and overlap between DG cores are allowed for at most one arm, but not both. Pseudo random numbers were generated with seeds to ensure reproducibility. (C-D) For each simulated DG dataset and clustering parameter combination (cluster, corelen, t_o and t_eig), the sensitivity and specificity of DG assembly was calculated for each of the top 100 DGs. For the 100 simulated DGs, the sensitivity of DG assembly is defined as the fraction of remaining alignments in each DG after CRSSANT DG assembly. For the top 100 CRSSANT assembled DGs, the specificity is defined as the fraction of alignments from the dominant simulated DG.

**Supplemental Figure 5.**
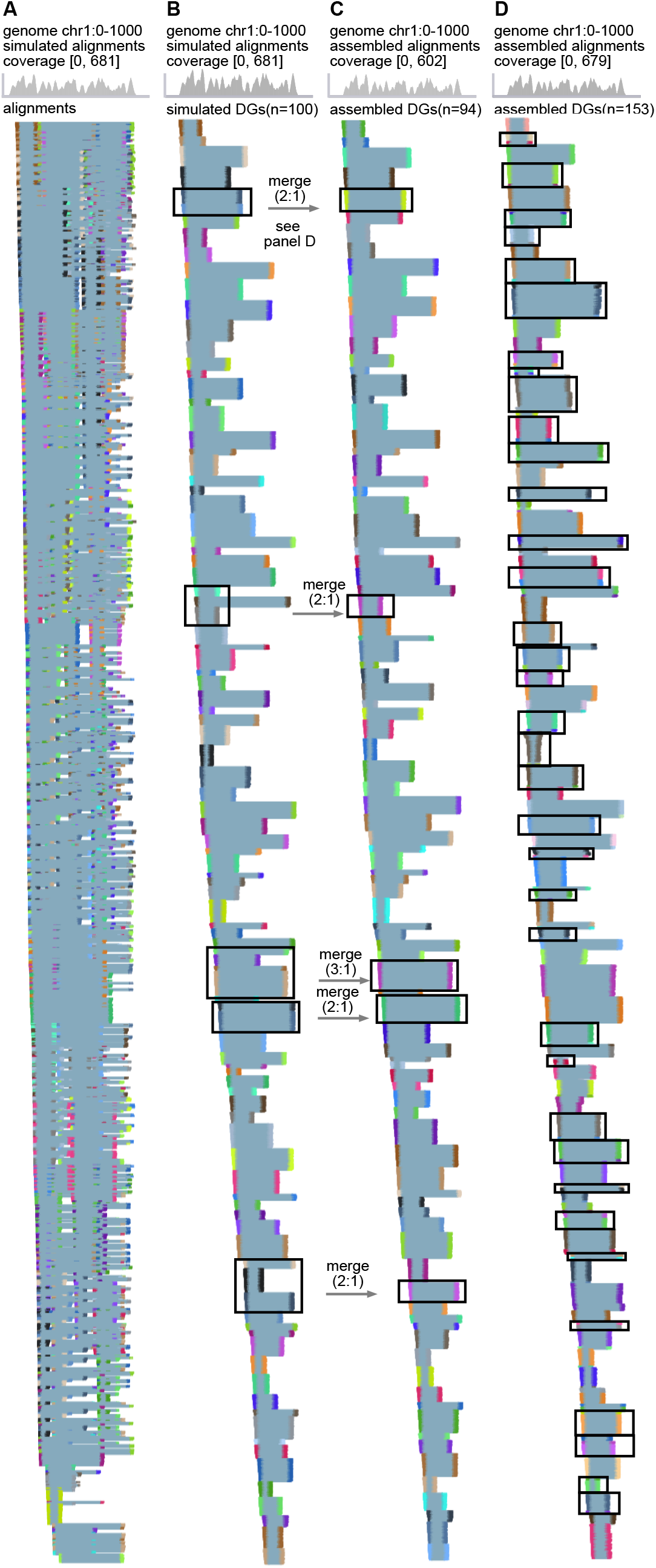
Comparison between simulated and CRSSANT assembled DGs. (A-B) Simulated 5335 alignments on chr1:1-1000 with the following parameters: coregap=50, corewithin=1000, edge in [5,15], numalign in [10,100], and numDGs=100. The alignments were shown without DG grouping (A) or grouped by the simulated DGs (B). In panel B, DGs were ranked based on the core start position from left to right. (C) The alignments in panel A were assembled using CRSSANT and the following parameters: threads n=8, cluster=clique, t_o=0.5 and covlimit=1000. Out of the 100 DGs, 5 merging events occured during CRSSANT assembly due to overlap of the two arms separately, one of which was merging 3 simulated DGs into one, therefore generating 94 assembled DGs. Otherwise, the DGs were highly consistent. (D) The alignments in panel A were assembled using CRSSANT and the following parameters: threads n=8, cluster=spectral, t_o=0.5, t_eig=5 and covlimit=1000. Out of the 100 DGs, ~ 50 of them were split into new DGs, even though they have significant overlaps. Example splitting events were highlighted with black boxes (compare to panel B). In addtition there are also a few merging events.

**Supplemental Figure 6.**
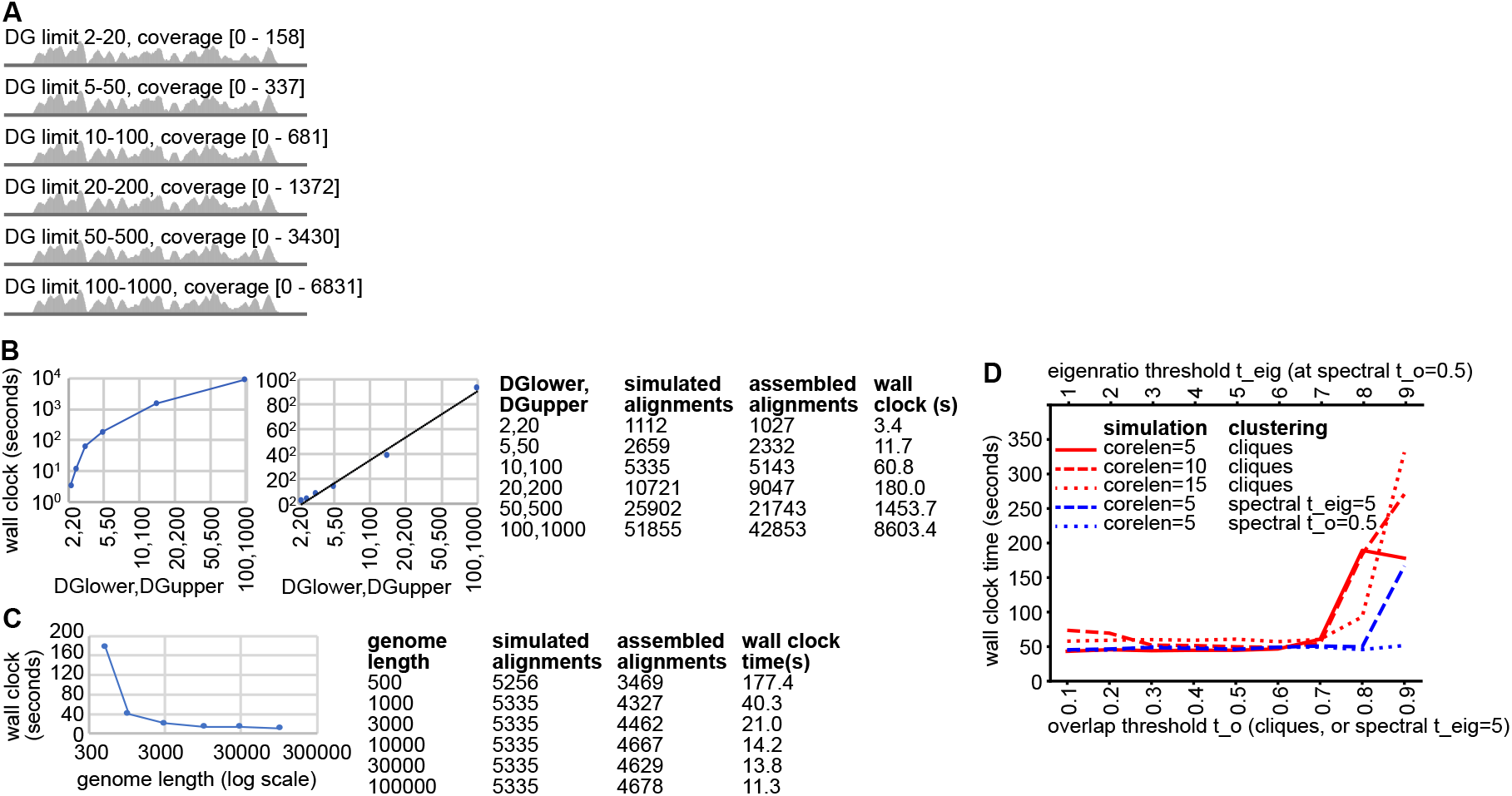
Benchmarking CRSSANT running time. (A) Six DG datasets were simulated using the following parameters: corelen=5, coregap=50, corewithin= 1000, edge in [5,15], numDGs=100. DG limit (DGlower and DGupper) were varied at the indicated values. (B) CRSSANT assembly of the various simulated DG datasets shown in panel (A) was performed using the following parameters: threads n=8, cluster=clique, t_o=0.5 and covlimit=1000. Running time were plotted in log (left plot) or square root (right plot) scales. (C) DG datasets were simulated with the default parameters, except genome length was varied between 300 and 100000. Then CRSSANT assembly of the datasets was run with the following parameters: threads n=8, cluster=clique, t_o=0.5 and covlimit=1000. Genome length was plotted in log scale. (D) Benchmarking the CRSSANT running time on 5335 alignments, with coregap=50, corewithin=1000, edge in [5,15], numalign in [10,100], and numDGs=100. Three simulated datasets were made with corelen=5, 10 or 15. Both cliques and spectral clustering methods were tested, at the specified parameters. The tests were run with threads n=8, covlimit=1000. All tests in this figure were run on a MacBook Pro (13-inch, 2017, Four Thunderbolt 3 Ports), 3.5 GHz Dual-Core Intel Core i7 processor, and 16 GB 2133 MHz LPDDR3 memory.

**Supplemental Figure 7.**
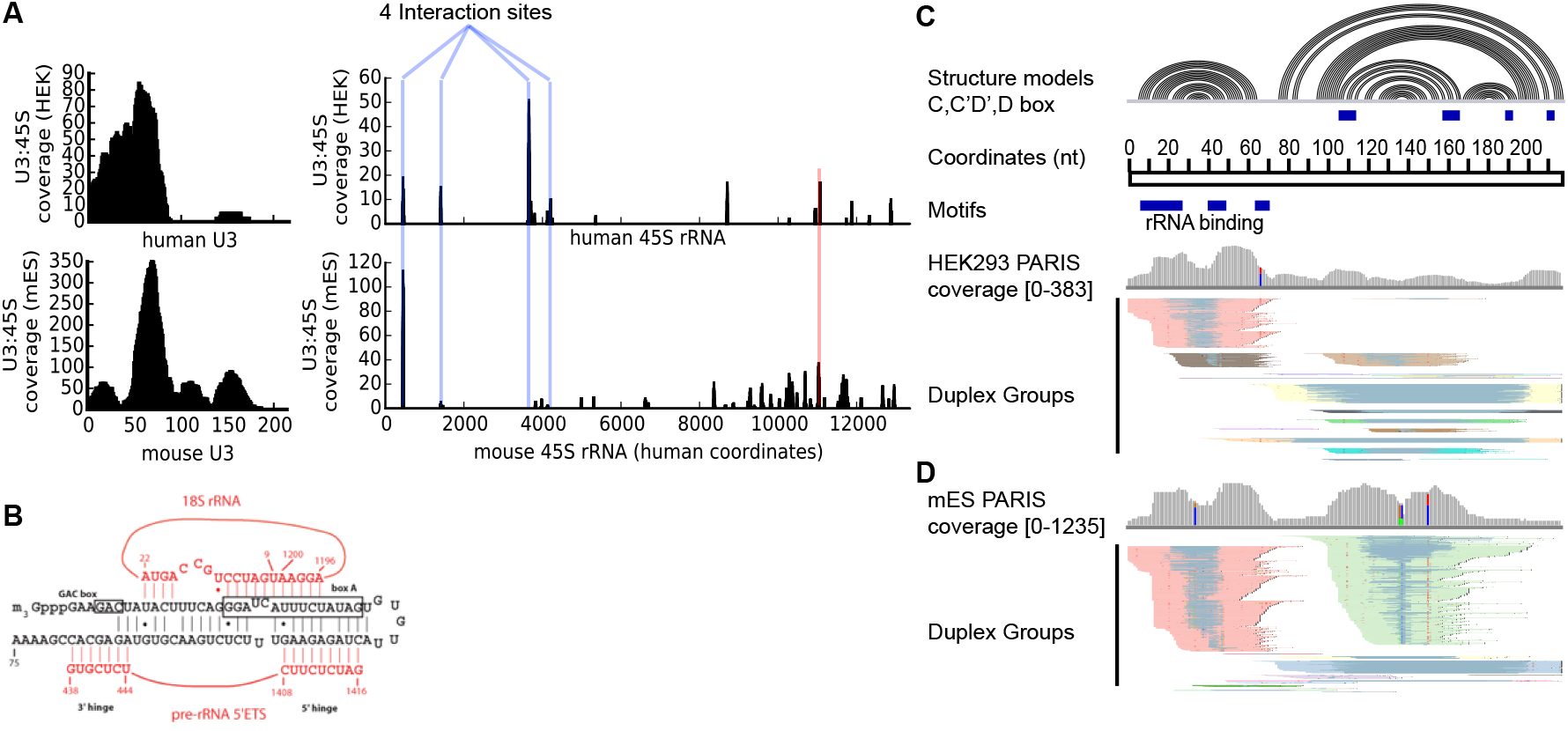
CRSSANT analysis of U3 snoRNA. The snoRNA U3 plays essential roles in ribosomal RNA processing through multi-partite interactions with the 45S rRNA precursor. (A) CRSSANT analysis of PARIS reveals 4 U3 binding sites on the rRNA precursor (blue vertical lines), using PARIS HEK293 cell data and mouse ES data (Lu et al. 2016), consistent with previous studies (Marz and Stadler 2009; Dutca et al. 2011). The red vertical line indicates a potential new interaction. (B) Previously published U3:rRNA interaction model, showing the 4 regions in 45S rRNA (red letters) that interact with the 5’ domain of U3 (black letters). U3:rRNA interaction model was from the snoRNA database (https://www-snorna.biotoul.fr/plus.php?id=U3). (C-D) CRSSANT analysis of human and mouse PARIS data support the proposed U3 secondary structures. The colors of the DGs are automatically determined by IGV and do not represent correspondence between human and mouse PARIS data or CRSSANT analysis. The 5’ end of U3 adopts a stem-loop conformation, in competition with the U3:rRNA interactions, similar to the U2 alternative conformations that mask the target recognition sequence. Very few low abundance DGs are inconsistent with the structure model (panels C-D, bottom). These low abundance DGs are likely the result of U3 dynamic folding intermediates or technical artifacts of crosslinking and proximity ligation. Together, these results provide further support for the CRSSANT method.

**Supplemental Figure 8.**
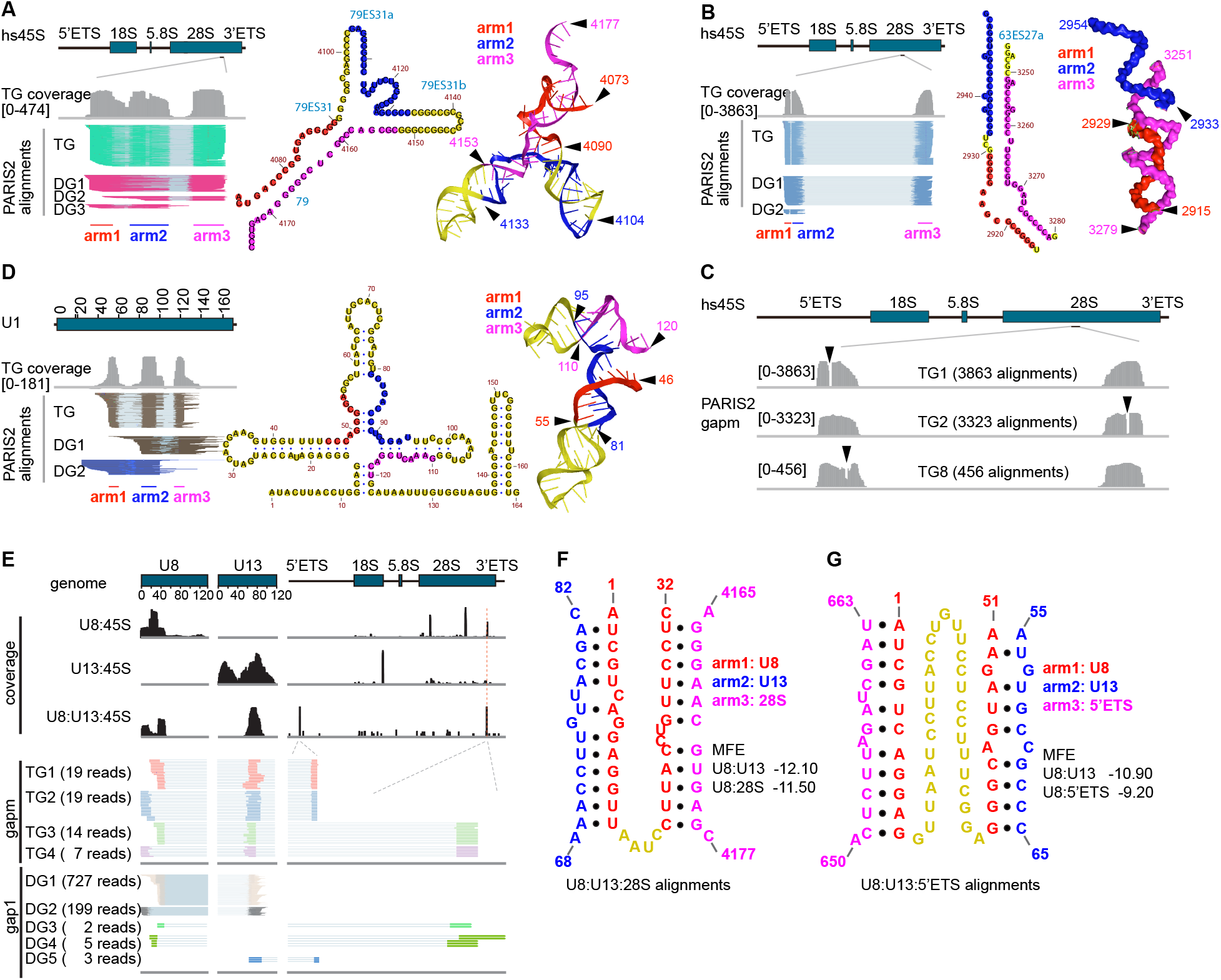
Gapm alignments and complex structures. (A) Gapm alignments supporting the two stemloops at a 4-way junction in the 28S rRNA. Left: alignments. Middle: secondary structure. Right: cryo-EM structure (PDB: 4V6X). (B) Gapm alignments supporting a longer dsRNA structure. The expansion segment 63ES27a cannot be resolved at high resolution by the cryo-EM, therefore the 3D model is only a fuzzy representation. (C) Most of the gapm alignments mapped to 28S rRNA come from longer dsRNA structures. In the top 10 TGs, 3 of them, shown here, came from the same longer dsRNA structure 63ES27a (3863+3323+456)/36689 = 21%). The first one is the same as in panel B. The RNase cleavage positions are indicated by arrowheads. (D) Gapm alignments showing two stemloops at the 4-way junction of the U1 snRNA. Left: alignments. Middle: secondary structure. Right: cryo-EM structure. (E) Gapm alignments showing mouse U8:U13:45S rRNA intermolecular interactions. The genome coverage of U8:mm45S (mouse 45S) and U13:mm45S rRNA interactions were presented separately. Colored alignments show the top 4 TGs and the DGs that compose the TGs, supporting U8:U13:mm45S rRNA interactions. All TG alignments were presented, while only 1000 alignments were used to prepare the DGs due to the much higher numbers of alignments for DGs. The U8:U13:mm45S interacting peak on mm45S was 20nt upsteam of the interacting peak of U8:mm45S. PARIS1 mES data were used here for this analysis. (F-G) Predicated basing pairing models of the two U8:U13:mm45S interactions.

**Supplemental Figure 9.**
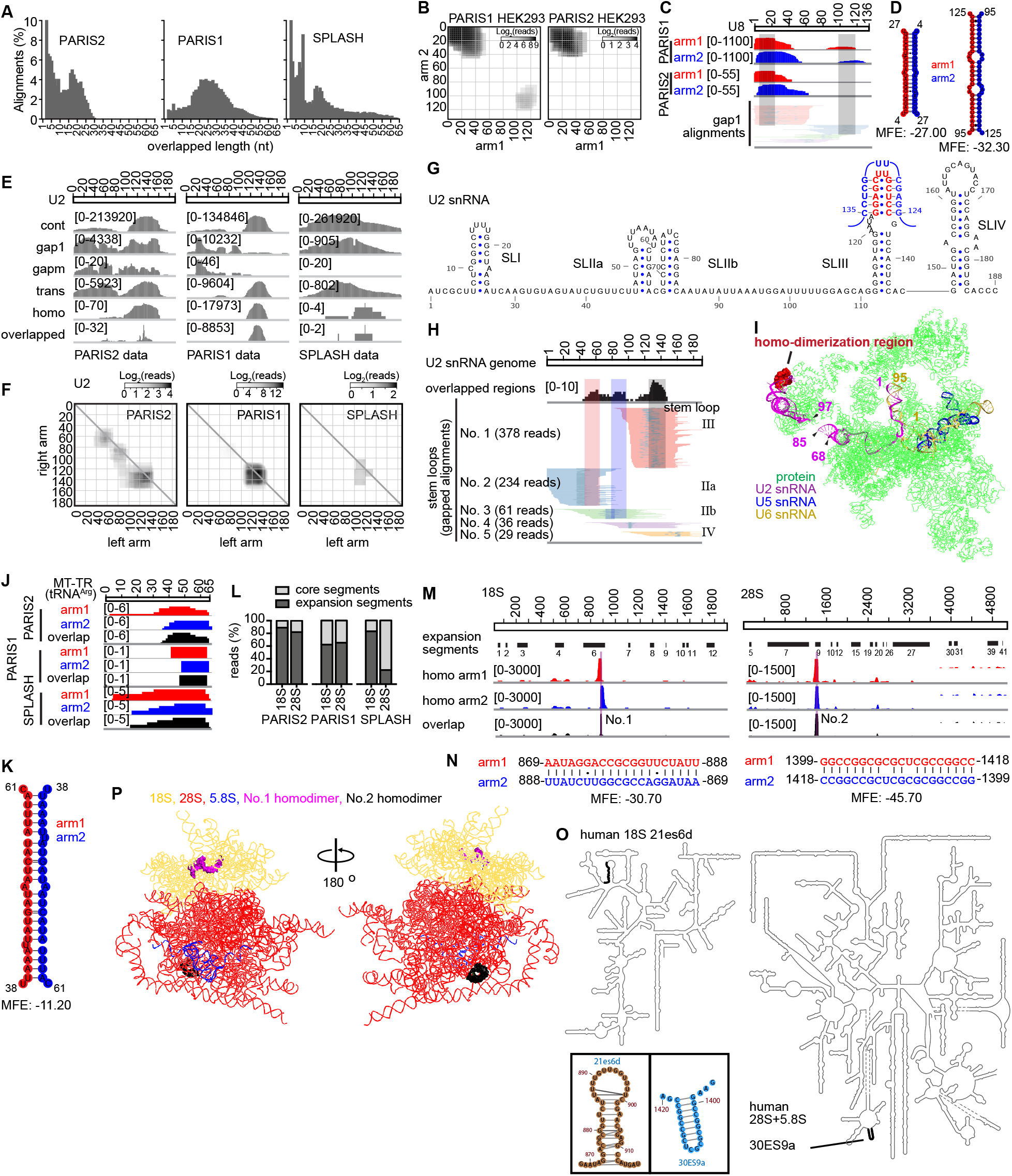
Identification of RNA homodimers in cellular RNAs. (A) Length distributions of overlapped regions in homo alignments in 3 datasets. Alignments with 1-2 nt overlaps were removed. (B) Heatmap of the U8 snoRNA homodimers identified in PARIS1 and PARIS2. PARIS1 and PARIS2 coverage was shown in log scale. (C) Correlation of U8 homodimers with local stem loops. PARIS2 HEK293 gap1 alignments were used to shown the local stem loops. (D) Predicated baseparing models of two U8 snoRNA homodimers. (E) Coverage of 5 different types of alignments on U2 snRNA from 3 different datasets. The overlapped region of homo alignments was shown separately at the bottom of each panel. (F) Heatmap of U2 snRNA homo alignment arms, with coverage shown in log scale. (G) 2D structure of U2 homodimer region on SLIII (blue and red letters). (H) Comparison of U2 overlapped regions with local stem loop DGs in the PARIS2 data. Only 1000 reads were used to assemble the DGs. (I) Physical location of U2 homo dimerization region on U2/U5/U6 snRNP cryo-EM structure (PDB: 7ABI). (J) A potential homodimer of the mitochondrial tRNA_Arg (chrM:10405-10469) based on overlapping alignments from 3 datasets. The top ranked homodimer regions in both rRNAs overlap expansion segments. (K) Secondary structure model of the MT-TR homodimer. (L) Distributions of homo alignments on core and expansion segments of 18S/28S rRNAs. (M) The PARIS2 identified the top2 rRNA homodimers, both of which are located in expansion segments. (N) Predicated base paring models of top 2 rRNA homo dimers. (O) Locations of the top 2 rRNA homodimers on the secondary structure models. The local stemloops were magnified and shown in the insets. (P) Locations of the top 2 rRNA homodimers on human ribosome cryo-EM structure (PDB: 4V6X), showing only the RNA chains.

**Supplemental Figure 10.**
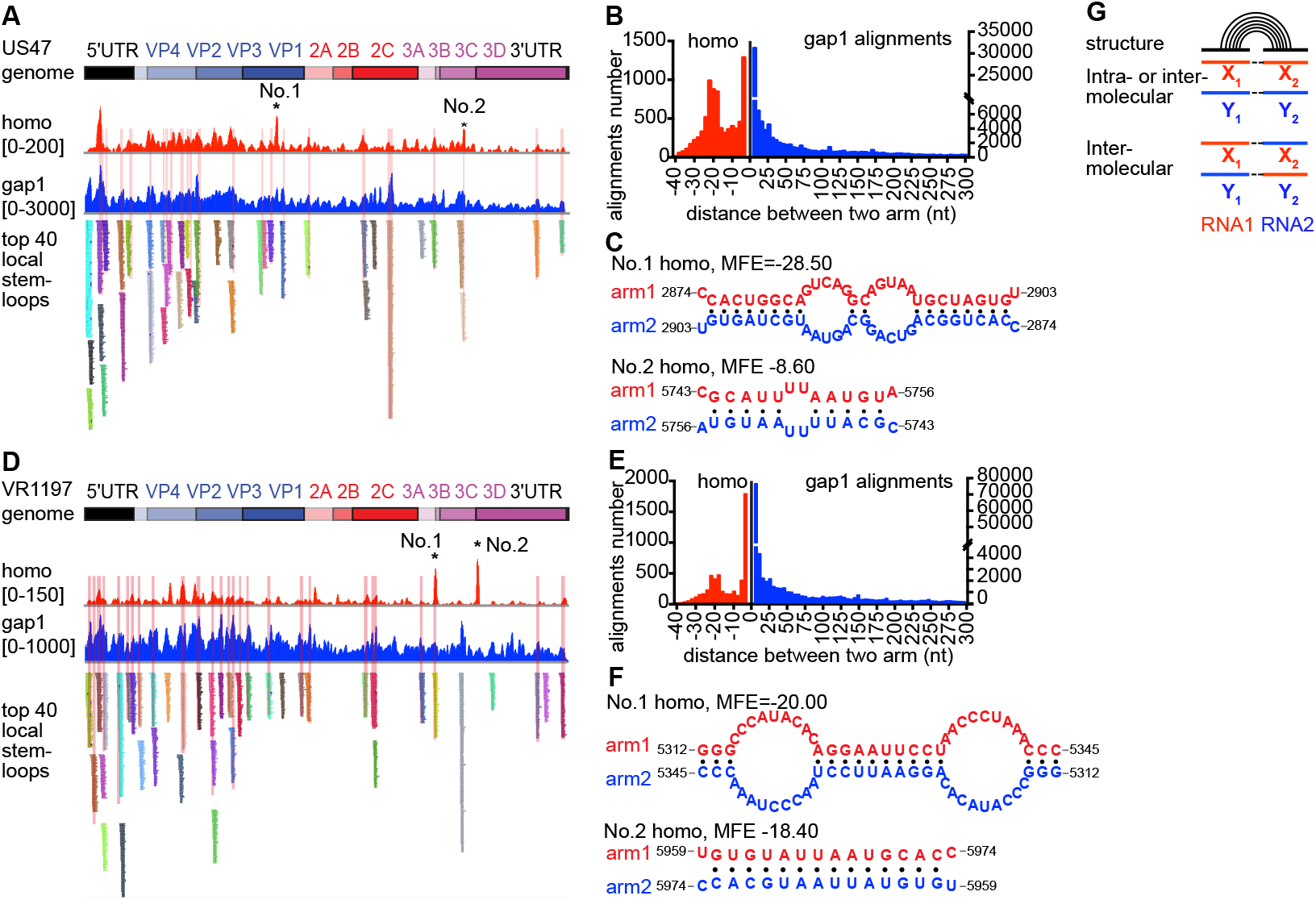
Analysis of potential homodimers in Picornavirus genomes. (A) PARIS2 overlapping alignments on US47 viral RNA (US/MO/14-18947). Gap1 alignments were shown as background. There two regions are prefer to form homodimer. Only homodimers with more than 2nt overlapping between two arms were shown here. The top 40 local stemloops were compared to the peaks of overlapping alignments. (B) The length distribution of distance between two arms. (C) The predicated base pairing models of No.1 and No.2 homodimer on the US47 viral RNA. (D) PARIS2 overlapping alignments on VR1197 viral RNA (F02-3607 Corn). Gapped alignments shown as background. Only alignments with more than 2nt overlapping between two arms were shown here. The top 40 local stemloops were compared to the peaks of overlapping alignments. (E) The length distribution of distance between two arms. (F) The predicated base pairing models of No.1 and No.2 homodimer on the VR1197 viral RNA. (G) An alternative method for the de novo identification of RNA homodimers. When RNA molecules from two different genetic backgrounds (red and blue lines) exist in the same cell, nucleotide sequence variations allow us to accurately map the fragments to the RNA of origin. When the two fragments are derived from the same genetic origin, the duplex could be either intra- or intermolecular. However, if the two fragments are from two different genetic backgrounds, then the duplex should be intermolecular.

**Supplemental Table 6.** List of top ranked RNAs with high homo alignment ratios. Homo alignments detected in three psoralen crosslinked sequencing data sets. RNAs were ranked by homo/gapped alignments ratio. Total alignements of each data are PARIS1 (GSE74353, HEK293 cells, 180,281,149 reads), PARIS2 (GSE149493, HEK293 cells: 84,612,226 reads; HeLa-US47: 92,779,821 reads; HeLa-VR1197: 71,775,448 reads) and SPLASH (SRX1715494 and SRX1715495, 104,655,285 reads).

**Top20 RNA homodimers in PARIS1 HEK293 data**
| RNAs | homo alignments | gapped alignments | homo/gapped ratio (%) |
| --- | --- | --- | --- |
| 28S | 26,592 | 10,614,406 | 0.25% |
| 18S | 12,807 | 5,339,371 | 0.24% |
| U8 | 6,681 | 1,349 | 495.26% |
| U2 | 6,054 | 19,820 | 30.54% |
| U1 | 3,164 | 274,878 | 1.15% |
| 5.8S | 1,107 | 894,490 | 0.12% |
| U6 | 120 | 35,324 | 0.34% |
| RN7SK | 104 | 100,378 | 0.10% |
| RN7SL | 72 | 89,613 | 0.08% |
| XIST | 23 | 54,187 | 0.04% |
| tRNAs | 20 | 177,947 | 0.01% |
| hs5S | 18 | 56,397 | 0.03% |
| U3 | 15 | 4,600 | 0.33% |
| RPPH1 | 14 | 7,073 | 0.20% |
| MT-CO1 | 10 | 19,215 | 0.05% |
| 16S | 10 | 30,765 | 0.03% |
| CARM1 | 8 | 598 | 1.34% |
| FRMD4B | 8 | 867 | 0.92% |
| TMEM178B | 8 | 1,340 | 0.60% |
| SON | 8 | 2,514 | 0.32% |

**Top20 RNA homodimers in PARIS2 HEK293 data**
| RNAs | homo alignments | gapped alignments | homo/gapped ratio (%) |
| --- | --- | --- | --- |
| 28S | 5,436 | 1,875,812 | 0.29% |
| 18S | 4,552 | 376,346 | 1.21% |
| U1 | 134 | 34,293 | 0.39% |
| 5.8S | 133 | 64,391 | 0.21% |
| U8 | 113 | 392 | 28.83% |
| RPL29 | 23 | 101 | 22.77% |
| 16S | 17 | 8,865 | 0.19% |
| tRNAs | 14 | 102,083 | 0.01% |
| RN7SK | 13 | 9,186 | 0.14% |
| U2 | 13 | 7,147 | 0.18% |
| 12S | 11 | 3,577 | 0.31% |
| RN7SL | 8 | 10,189 | 0.08% |
| MT-CO1 | 7 | 2,375 | 0.29% |
| MT-CO2 | 7 | 1,739 | 0.40% |
| XIST | 7 | 3,488 | 0.20% |
| CPB1 | 5 | 2 | 250.00% |
| MT-ATP6 | 5 | 1,307 | 0.38% |
| MT-ND5 | 5 | 784 | 0.64% |
| MT-TR | 5 | 144 | 3.47% |
| MT-ND4 | 4 | 871 | 0.46% |

**Top13 RNA homodimers in SPLASH data**
| RNAs | homo alignments | gapped alignments | homo/gapped ratio (%) |
| --- | --- | --- | --- |
| 5.8S | 12,003 | 165,510 | 7.25% |
| 28S | 5,827 | 634,643 | 0.92% |
| 18S | 5,641 | 491,035 | 1.15% |
| U1 | 1,076 | 81,758 | 1.32% |
| BET1L | 9 | 2 | 450.00% |
| RN7SL | 8 | 3,218 | 0.25% |
| COX7A2L | 4 | 6 | 66.67% |
| POLR2J | 3 | 4 | 75.00% |
| ADAR | 2 | 12 | 16.67% |
| LONP1 | 2 | 1 | 200.00% |
| NCOR2 | 2 | 7 | 28.57% |
| U8 | 2 | 224 | 0.89% |
| 12S | 1 | 489 | 0.20% |

| RNAs | homo alignments | gapped alignments | homo/gapped ratio (%) |
| --- | --- | --- | --- |
| US47 | 9,874 | 922,045 | 1.07% |
| VR1197 | 4,648 | 442,163 | 1.05% |

